# The Effect of Resistance Training Volume on Individual-Level Skeletal Muscle Adaptations: *A Novel Replicated Within-Participant Unilateral Trial*

**DOI:** 10.1101/2025.07.24.666533

**Authors:** Zac P. Robinson, James Steele, Eric R. Helms, Eric T. Trexler, Michael E. Hall, Chun-Jung Huang, Joshua C Pelland, Jacob F Remmert, Seth R. Hinson, Scott A. Mikula, Alexandre A. Hamaïde, Michael C. Zourdos

## Abstract

**Purpose:** There is growing emphasis on investigating heterogeneity in resistance training (RT) outcomes, likely motivated by observations of substantial gross variability in training effects. However, gross variability does not necessarily represent true inter-individual response variation (IRV) and can be obscured by measurement error, sampling variance, and biological variability. Appropriate study design and statistical analysis are required to distinguish IRV from these confounding sources of within-participant variation.

**Methods:** 16 recreationally trained participants completed a novel replicated within-participant unilateral design across two 11-week training phases separated by a 6-8 week washout. Lower limbs were randomized to a low volume (∼8 sets/week) or high volume (∼16 sets/week) training protocol in each phase. We assessed both general (GEN; average response across conditions) and condition-specific (CON; difference between volumes) IRV for vastus lateralis cross-sectional area and leg press one-repetition maximum using a multi-stage statistical approach.

**Results:** Higher weekly set volumes demonstrated a detectable advantage for muscle hypertrophy (1.8 cm² [95% HDI: 0.29, 3.41]; 98.77% posterior probability) but not maximal strength (3.48 kg [95% HDI: -5.1, 12.15]; 80.01% posterior probability). Despite substantial gross variability, we failed to detect irrefutable evidence of meaningful IRV. Integrated methods revealed stronger evidence for GEN versus CON IRV, with correlation coefficients ranging from 0.67 to 0.7 for GEN versus 0.04 to 0.06 for CON.

**Conclusions:** Our findings clearly illustrate that gross variability in training outcomes does not necessarily indicate true inter-individual differences, a distinction critical for both research and practice.

**KEY POINTS:**

- Gross variability in resistance training outcomes is commonly interpreted as evidence of meaningful inter-individual response variation (IRV), but this variability often reflects confounding sources of within-participant variation rather than true inter-individual differences.
- Using a novel replicated within-participant unilateral design, we investigated the effects of different weekly set volumes on changes in muscle size and maximal strength, as well as the inter-individual variation thereof.
- At the group level, higher weekly set volumes produced modest but detectable benefits for muscle hypertrophy but not maximal strength. At the individual level, we failed to reveal irrefutable evidence of true IRV for either primary outcome, though there was stronger support for variability in training responses independent of weekly set volume (i.e., general IRV) compared to differential responses between volume conditions (i.e., condition-specific IRV).
- While this study has limitations, it highlights that appropriate study design and statistical analysis are essential for investigating IRV in resistance training. Our findings, along with the broader research in multiple disciplines (i.e., medicine, nutrition science, and exercise physiology), demonstrate that gross variability can purely reflect confounding sources of within-participant variation (e.g., sampling variance, measurement error, biological variability) rather than true inter-individual differences—a distinction critical for both research and practice.

## INTRODUCTION

The pursuit of nearly all scientific inquiry is to explain or predict variability. In disciplines such as medicine, nutrition science, and exercise physiology, there is growing emphasis on quantifying the variability in the effect of an intervention (i.e., variance) rather than focusing solely on the expected magnitude of effect (i.e., mean) (Senn, 2015; Robinson *et al*., 2024a; Lolli *et al*., 2025). This shift reflects the desire to provide precise, individualized recommendations that theoretically yield superior outcomes compared to population-based approaches. Indeed, many studies have reported a high degree of *gross variability* (i.e., wide range of change scores following an intervention) that would seemingly warrant highly personalized recommendations (Hubal *et al*., 2005; Erskine *et al*., 2010; Ahtiainen *et al*., 2016; Damas *et al*., 2019; Hammarström *et al*., 2020; Lixandrão *et al*., 2024). However, recent evidence indicates that gross variability alone does not represent *true* inter-individual response variation (IRV) (Atkinson & Batterham, 2015; Robinson *et al*., 2024a). This distinction arises from the challenge of delineating the variation of interest from other confounding sources of within-participant variation (WPV), such as sampling variance, biological variability, and measurement error. Accurately identifying true IRV requires appropriate pairing of study design and statistical analysis.

One approach to partially detect IRV involves comparing the variability of a control condition to that of a condition undergoing an intervention. This method isolates the *intervention-induced variability*, within which true IRV may exist. Multiple studies across the aforementioned domains applying this approach have found that the variability introduced by an intervention ranges from modest to undetectable, contrasting expectations based on gross variability (Williamson *et al*., 2018; Kelley *et al*., 2021, 2023, 2024; Esteves *et al*., 2021; Kelley *et al*., 2022; Bonafiglia *et al*., 2022). In the context of resistance training (RT), Steele et al. found that non-training controls exhibited *greater* variability in strength and hypertrophy outcomes than RT conditions after accounting for the inherently positive relationship between the mean and variance in biological systems (i.e., Taylor’s law) (Taylor, 1961; Steele *et al*., 2023). This finding suggests that variation in RT outcomes is largely explained by sources outside of true inter-individual differences. However, this approach has multiple limitations. First, non-training control conditions necessitate the use of untrained individuals, which may limit the generalizability of these findings to individuals with resistance training experience. Second, this approach focuses on *general* responses to RT rather than examining the variability between *specific protocols or conditions* (e.g., high versus low weekly set volume). This omission is crucial, as hypotheses regarding inter-individual differences in the protocols or conditions that optimize training outcomes are widespread in both research and practice.

To most effectively investigate IRV, replication of an intervention within the same cohort of individuals is likely necessary (Senn *et al*., 2010; Senn, 2024). Repeated exposure to the intervention allows for estimation of the *magnitude* (i.e., quantifying the expected effect of the intervention and precision thereof), *reliability* (i.e., correlation between replicates of the interventions), and *agreement* (i.e., differences between replicates of the interventions) of individual training effects. These estimates explicitly partition the sources of training-induced variation from WPV, thereby determining whether: i) true IRV is detectable (i.e., sufficient correlation between replicates of the interventions exists) and ii) the magnitude of this variation is meaningful (i.e., differences between individuals exceed differences between replicates of the interventions for a given individual). These estimates can be derived using both naïve and integrated statistical methods, with the former calculating individual effects using only participant-specific data, while the latter incorporates information from the implied population (i.e., sample) to enhance the precision of estimates for a given individual (e.g., partial pooling and shrinkage) (Araujo *et al*., 2016).

Therefore, this study aimed to investigate both general (GEN) and condition-specific (CON) IRV using a novel replicated within-participant unilateral design in trained participants. Although exploratory in nature, we hypothesized that, despite observing clear gross variability, there would be no clear evidence of true IRV for the primary outcomes. Additionally, we hypothesized that muscle hypertrophy would be enhanced by higher weekly set volumes on average, while strength gains would not.

## METHODS

### Ethical Approval

The methods, hypotheses, and planned analyses for this study were originally pre-registered (https://osf.io/aw5zx). However, throughout the data collection period (but prior to examining any data), our hypotheses and views on how to best communicate, analyze, and report our findings changed substantially due to improved understanding of the topic. Thus, we deviated from some specifications in the pre-registration and evaluated the outcomes in an exploratory fashion. All individuals provided written consent prior to participation and the University’s Institutional Review Board approved this investigation (IRBNET ID: 1955508-5).

### Sample Size Justification

Given the resource-intensive nature of the current study design, our sample size is justified by feasibility (Lakens, 2022). As many participants as possible were recruited throughout the data collection period determined by the lead author’s doctoral committee, but due to the limited sample size, the data are presented such that they can be meta-analytically aggregated in the future.

### Participants

A mixed-sex sample of 16 recreationally trained individuals (≥2 consecutive years of RT experience with ≥ 2 sessions per week) participated in this study as determined by a training history questionnaire (Table 1). Participants with contraindications to exercise as determined by a health history questionnaire (i.e., heart disease, hypertension, diabetes, etc.) were excluded. Approximately one week prior to participation in each phase, all participants were asked to cease all supplementation (other than habitual caffeine intake) and external recovery practices (e.g., foam rolling) for the remainder of the intervention period.

**Table 1.** Participant baseline characteristics. Data are mean ± standard deviation.

| Participant Baseline Characteristics |  |  |
| --- | --- | --- |
| Metric | Male | Female |
| n | 11 | 5 |
| Age (years) | 22.40 ± 2.07 | 21.40 ± 2.41 |
| Training Exp. (years) | 5.33 ± 2.94 | 3.83 ± 1.60 |
| Body Mass (kg) | 87.92 ± 14.34 | 67.12 ± 8.69 |
| Height (cm) | 175.65 ± 7.06 | 163.00 ± 5.68 |
| Knee Flexion (°) | 107.50 ± 6.45 | 105.00 ± 5.00 |
| Hip Flexion (°) | 125.29 ± 10.98 | 128.75 ± 8.54 |
| Skinfold Sum (mm) | 39.23 ± 16.13 | 57.00 ± 13.92 |
| Data are Mean ± SD. |  |  |

The post-testing measurements of the second training phase were not completed for two participants (n=1 dropout and n=1 training-related injury). Additionally, one participant could not perform isometric strength testing at 2 of the 3 joint angles without discomfort; thus, only data from the asymptomatic joint angle was used for this participant. Additional details regarding the participants’ habitual training style and training performed throughout the intervention period can be found in the supplementary materials (https://osf.io/rghe4).

### Experimental Design

This study utilized a replicated within-participant unilateral design that lasted 28-30 weeks and consisted of training phase 1 (11 weeks), a washout period (6-8 weeks), and training phase 2 (11 weeks). Lower limbs were initially randomized to either a low (LV = 8 sets per week) or high volume (HV = 16 sets per week) training condition for the initial 11-week phase. The weekly set volumes were performed with the unilateral leg press evenly split over two training sessions separated by approximately 72 hours (e.g., Monday and Thursday). Participants’ dominant limbs were counterbalanced so there were an equal number of dominant limbs assigned to the two training conditions at baseline.

Following a 6-8 week washout period, in which participants were asked to return to habitual training, participants performed a second 11-week phase that was identical to the first except that each participant’s limbs were re-randomized to the training conditions (i.e., either HV or LV). This re-randomization creates four possible sequences across the entire intervention where each phase represents an independent randomized trial. If participants were instead required to deterministically switch conditions in the second phase, the condition allocation would be predetermined based on the first assignment, which would violate the principle of independent replication.

During each 11-week phase, week 1 served as initial baseline testing of outcome measures, while weeks 2 and 3 served as an introductory period in which both limbs performed lower-volume training (i.e., LV = 6 sets per week, HV = 12 sets per week) in preparation for the main training program and to ensure both limbs had a similar relative increase in weekly set volume heading into the main training program (i.e., ∼25% from peak volume). In week 4, participants repeated the testing procedures from week 1 and completed two “mock” training sessions (one on each limb) that were identical to the main training program (i.e., 4 and 8 sets per session) to gradually introduce them to the volumes of the main training program. Weeks 5-10 served as the main training program with post-study testing in week 11. A timeline of the procedures is in Figure 1.

**Figure 1:**
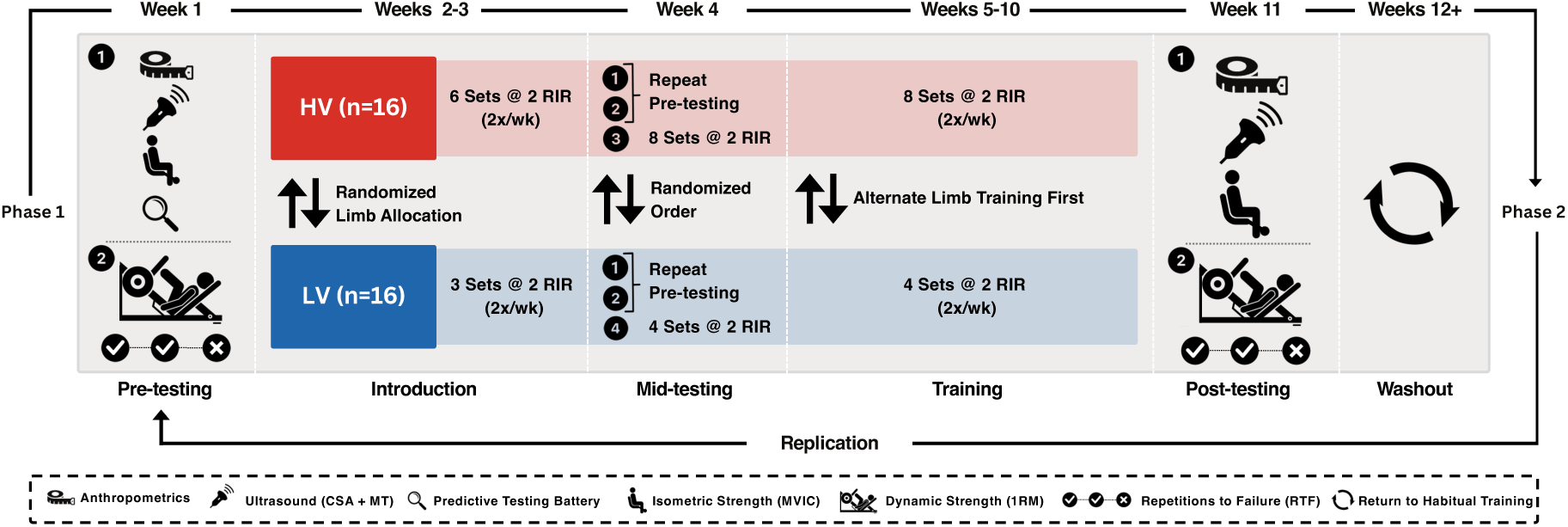
Summary of study design

At baseline in each phase, anthropometrics were assessed as were various metrics (e.g., vertical jump, estimated maximal oxygen consumption, etc.) that we planned to explore as potential predictors of IRV if sufficient evidence to warrant this step was identified (https://osf.io/wfk7x). Additionally, subjective scales assessing limb-specific muscle soreness and session rating of perceived exertion (sRPE) were administered immediately before and after each training session, respectively.

The primary outcomes of this study were vastus lateralis (VL) cross-sectional area (CSA) and unilateral leg press one-repetition maximum (1RM), while VL muscle thickness (MT) and peak force from a unilateral knee extension maximal voluntary isometric contraction (MVIC) were considered secondary outcomes. Each of these outcomes was assessed during week 1 (baseline), week 4 (midpoint), and week 11 (post) in each phase. A timeline of the entire study including when each measurement was assessed can be seen in Figure 1.

### Testing Procedures

#### Resistance Training Program

In both phases, participants trained the unilateral leg press (Leverage Horizontal Leg Press, Body-Solid, Forest Park, IL, USA) twice per week. One limb performed LV training which consisted of 6 sets per week in weeks 2-3 and 8 sets per week in weeks 4-10, while the other limb performed HV training which consisted of 12 sets per week in weeks 2-3 and 16 sets per week in weeks 4-10. There were 3 minutes of rest between sets and all sets were completed on one limb prior to beginning sets on the other limb. The limb that was trained first in the session was alternated every session. Prior to each training session, participants completed a dynamic warm-up followed by a specific warm-up on the unilateral leg press. The specific warm-up was: no load for as many repetitions as desired, 40% of the working load for 8 repetitions, 75% of the working load for 6 repetitions and finally 85% of the working load for 4 repetitions.

During weeks 4-10 in both study phases, the target training load was 8-12 repetitions per set with 2 repetitions in reserve (RIR) or what equates to a 10-14 RM. To objectively determine the set endpoint at 2 RIR, participants performed 4 sets to momentary failure with the newly established 1RM at weeks 1, 4, and 11 in each phase and mean propulsive velocity (MPV) was assessed throughout the set (Linear Encoder, Vitruve, Madrid, Spain). The 4 sets were averaged and the MPV range associated with each RIR was determined (Pelland *et al*., 2022). Thus, sets in weeks 4-10 were then terminated upon MPV initially reaching that range. To illustrate, if across 4 sets the average MPV of the final 4 repetitions was 0.35, 0.30, 0.25, and 0.20 m·s^-1^ for a given participant, this would correspond to velocity ranges of greater than or equal to 0.35 m·s^-1^ for 3 RIR, 0.30-0.34 m·s^-1^ for 2 RIR, 0.25-0.29 m·s^-1^ for 1 RIR, and less than or equal to 0.20 m·s^-1^ for 0 RIR. In this example, each set would have been terminated as soon as a repetition registered a velocity that indicated <3 RIR.

However, near the end of phase 2 data collection for the first 7 participants, the linear position transducer used to measure MPV began to malfunction. Thus, for the remainder of the study a subjective RIR termination method was utilized. Participants were instructed to terminate the set when they believed they had 2 RIR. The supervising researchers made it clear that if a participant was unsure if the proximity to failure was sufficient, to always err on the side of terminating the set too close to failure. For all sets throughout the study (i.e., even those with velocity-based termination) subjective RIR was also recorded.

The load for the first training session of both phases was set to 75% of 1RM for both limbs. Load was progressed for the following set by 3% per repetition over the target if the sum of the repetitions performed and RIR was >14, while load was decreased by 6% per repetition under the target if the sum was <10. The average load of the previous session was used as the initial set load in each session. After completing the final testing week of the first training phase, participants entered a 6-8 week washout period in which they were instructed to return to their habitual training style. The length of the washout period was individualized necessarily due to participant availability. The second phase of the intervention was identical to the first except each participant’s limbs were re-randomized into the training conditions.

#### Unilateral Leg Press Execution

The leg press settings and foot placement were adjusted for each individual so they achieved as much knee flexion as possible without their glutes coming off the seat or heels coming off the platform. On each repetition, in both testing and training sessions, participants paused in the bottom position until the load became motionless. Then, the investigator verbally indicated to the participant to perform the concentric portion of the repetition with maximal intended velocity. Throughout the intervention, some participants reported minor discomfort when performing the leg press. In these cases, minor adjustments to the leg press settings and foot placement were permitted but any changes were always mirrored by both limbs.

#### Additional Exercise

Following the primary portion of each training session (i.e., leg press) participants also performed the bilateral seated leg curl (Leg Extension Curl Machine, Titan Fitness, Memphis, TN, USA). This exercise was not experimentally manipulated, rather was performed to aid in participant recruitment and ensure participants were able to train all major muscle groups. The machine was adjusted so that each participant sat in 90 degrees of hip flexion and trained from 0-90 degrees of knee flexion. Participants completed 3 sets per session, aiming for 10 repetitions at a subjectively reported 2 RIR. Load was adjusted ∼2% per repetition or RIR off of the target. 3-minute rest periods were taken between sets. The upper body training of all participants was controlled as part of separate research projects in our laboratory. Finally, participants were permitted to perform cardiovascular exercise of a subjectively defined low to moderate intensity 2 days per week throughout the intervention. However, these sessions could not be within the 24 hours before a training session or the 48 hours before a testing session. Participants were asked to keep all habitual physical activity consistent between phases.

#### Unilateral Leg Press One-Repetition Maximum

All participants began strength testing with a 5-minute dynamic warm-up, followed by a leg press-specific warm-up where they performed as many repetitions as desired with an empty sled. Next, they executed a series of warm-up sets, starting with 5 repetitions at 20% of their estimated 1RM, then 3 repetitions at 50%, 2 repetitions at 70%, and finally 1 repetition at 80%. After completing the set with 80% of their estimated 1RM, participants rested for 3 minutes before a final warm-up lift using a weight between 85-90% of their estimated 1RM, as determined by the investigators. Throughout the warm-up if the participant requested to repeat a warm-up (i.e., if they did not feel sufficiently prepared for the next load), they were permitted to do so, but the additional warm-up set was also performed on the other limb.

Following the last warm-up set, there was a 3-minute rest period while the investigators set the initial weight for the first 1RM attempt. The weight was increased with each subsequent attempt to determine the participant’s 1RM, with a 3-minute rest interval between each lift. During each warm-up and 1RM attempt, the participant’s perceived RIR and MPV were collected to assist in determining the appropriate weights for the attempts. After completing 1RM testing on one limb, participants had a 5-minute rest before repeating the protocol on the opposite limb. All 1RM tests were conducted using Eleiko lifting discs (Eleiko, Austin, TX, USA), which are calibrated to the nearest 0.25 kg.

#### Repetition Performance Assessment

Each time 1RM was tested (weeks 1, 4, and 11 in each phase), participants completed a repetition performance test after achieving a 1RM on both limbs. Participants used 70% of the just-established limb-specific 1RM and performed 4 sets to momentary failure (the point at which participants could not complete another repetition despite maximal effort to do so) with strong encouragement from the investigators to reach the appropriate set end point. Between sets, participants rested 3 minutes with the limbs lying on the leg press in front of them in a standardized position. During each set, the MPV was collected for each repetition and the total number of repetitions performed. These sets were used to determine the MPV ranges for an objective 2 RIR set endpoint as described earlier.

#### Vastus Lateralis Cross-Sectional Area and Muscle Thickness

VL CSA and MT were evaluated using panoramic B-mode ultrasonography (iU22, Philips, Cambridge, MA) which has shown sufficient reliability to measure muscle size and changes thereof following resistance training (Valera-Calero *et al*., 2021). Initially, the investigator located the origin of the VL via ultrasound, marking the superior border roughly in alignment with the lateral epicondyle of the femur. A second mark was then placed midway between the superior border and the lateral epicondyle. The medial and lateral edges of the VL were identified via ultrasound at points perpendicular to this midpoint. Another mark was placed midway between these borders to establish the final scanning line. A piece of tape was used to connect these three points—the lateral epicondyle, midpoint of the VL, and the superior border—as a reference plane.

Scans were conducted along this tape in a longitudinal, controlled, and steady motion, maintaining the probe’s perpendicular orientation to the muscle while taking care not to deform the underlying tissue. A generous amount of water-soluble gel was applied to both the skin of the participant and the ultrasound probe to best achieve acoustic coupling. Two scans were captured at each timepoint. The resulting Digital Imaging and Communications in Medicine (DICOM) files were uploaded into ImageJ (National Institute of Health, Bethesda, MD, USA), where the scale was set. The vastus lateralis outline was then traced using the polygon tool to calculate the CSA. After tracing, the centroid of the polygon was determined, and a rectangular border was positioned around the entirety of the vastus lateralis polygon. The image was rotated to align the vastus lateralis borders at the centroid horizontally, facilitating a MT measurement at this point (https://osf.io/su7qg). To our knowledge, this method of MT assessment has not been validated and thus, was considered a secondary measure of muscle size. All scanning and analysis tasks were performed by the same investigator.

#### Anthropometrics

Total body mass (kg) was measured by a calibrated digital scale (Mettler-Toledo, Columbus, Ohio, USA) and participants’ height (cm) was measured via a wall-mounted stadiometer (SECA, Hamburg, Germany). A proxy of body composition was estimated using the average of two skinfold thickness measurements acquired from the sum of three sites: chest, abdomen, anterior thigh for males and triceps, suprailiac, and anterior thigh for females. If any measurement was >2mm different than the previous measure, a third thickness was taken. Additionally, the circumference of the thigh at 30, 50, and 70% of femur length was also recorded (measured along the same scanning plane used for the ultrasound measures). The same investigator took all measurements.

#### Unilateral Knee Extension Maximal Voluntary Isometric Contraction

Each participant performed two maximal isometric knee extension contractions per limb at three different joint angles determined by goniometer (110, 90, and 70 degrees of knee flexion) for a total of 6 maximal contractions each 3 seconds in duration. Attempts at each joint angle were alternated between limbs prior to moving to the next joint angle with a 2-minute rest between attempts and a 4-minute rest between attempts of the same limb. The tests were performed in a randomized order between limbs but kept consistent at each timepoint for each participant.

Testing was performed on a modified leg extension device (Titan Fitness, Item No. 401556) instrumented with a load cell (Honeywell Model 125 S-type/z-beam Load Cell, (06-K627-03), Charlotte, NC, USA) at a sampling rate of 1kHz. A custom-built mechanism was attached to the device that allowed for individual adjustments around the joint angle of the knee to measure peak isometric force. Given that our custom device has not been validated, this method was considered a secondary measure of maximal strength. To allow for joint multivariate modeling with the primary strength outcome (i.e., 1RM), the available measurements were averaged across joint angles and attempts such that each time point was represented by a single value per limb.

#### Subjective Proxies of Fatigue

Immediately before each training session, muscle soreness was recorded for the quadriceps and glutes of each individual limb with a custom scale adapted from Vickers (Vickers, 2001). The scale ranges from 0-6 with “0” indicating the participants had a complete absence of soreness and a score of “6” indicating a severe pain that limits their ability to move. Immediately following each training session, each participant recorded their limb-specific sRPE (Day *et al*., 2004). The scale ranges from 0-10 with “0” indicating the participant was at “rest” and a score of “10” indicating “maximal effort”.

### Inferential Overview

All analyses were performed in R (version 4.4.1, The R Foundation for Statistical Computing, 2025) and all data, code, and supplementary materials can be accessed at https://osf.io/g3vbe/. In addressing our research questions for the primary outcomes (i.e., CSA and 1RM), we took a systematic multistage approach. Secondary outcomes (i.e., MT and MVIC) were analyzed with identical procedures but in order to adequately report, interpret, and discuss the implications of the primary results, all visualizations of these secondary outcomes are included in the supplementary materials. Specifically, our analysis was separated into the following stages:

1. *Test-Retest Reliability*
2. *Gross Variability*
3. *Naïve Estimation of Inter-individual Response Variation*

*ii) Magnitude*
*iii) Reliability*
*iv) Agreement*
4. *Integrated Estimation of Inter-individual Response Variation*

*ii) Magnitude*
*iii) Reliability*
*iv) Agreement*
5. *Prediction*
6. *Simulation*

These stages make two important distinctions: i) the methods by which training effects are estimated (i.e., naïve or integrated through formal statistical modeling) and ii) the domain in which the estimates of IRV apply (i.e., GEN or CON). Naïve estimation refers to the calculation of effects without borrowing information from the sample/implied population participants exist within. Alternatively, integrated estimation formally incorporates this information in a statistical model and adjusts the results to potentially improve the accuracy and precision of the estimated parameters. Specifically in the context of IRV, “shrinkage” of observations towards the implied population mean may, potentially counterintuitively, offer superior estimates on the individual-level than when using the individual-level information alone (Araujo *et al*., 2016). Given our small sample size and the complexity of statistical models necessary to investigate our research questions (Senn, 2017), we leveraged both approaches to potentially detect sensitivity in our results to the analytical approach utilized; though we view the results from the integrated estimation with the most weight.

Second, the two specified domains (i.e., GEN and CON) refer to the average outcome across conditions (i.e., main effect of time) and the difference in outcomes between conditions (i.e., time-condition interaction effect), respectively. These domains characterize two target sources of variability: i) IRV in training effects independent of experimental condition, and ii) IRV within the experimental condition that results in a superior training effect. The distinction between these two domains will be made herein. Importantly, due to the lack of a time-matched non-training control group, we must make the assumption that changes in our primary outcomes are indeed caused by the training intervention rather than other confounding sources of error (e.g., measurement drift). While this is a limitation, the ability to obtain unbiased estimates of causal effects in the absence of a time-matched non-training control group seems to adequate in RT research (Steele *et al*., 2023).

Crossover study designs are not without critical assumptions. In the present study, we directly account for potential bias due to phase-relevant effects (i.e., main effect of phase, time-phase, and time-condition-phase interactions). However, our study design is not capable of causally separating carryover effects (i.e., the effect of the previous intervention influences the current intervention) from these interaction terms. Moreover, our target variance estimates assume consistency of the individual-level outcomes across phases (i.e., assumes a single true population variance to be estimated).

This assumption is necessary because each participant contributes a single observation for the time-phase and time-condition-phase interactions, and thus, the variances of these terms cannot be causally identified. To overcome this limitation, even more complex study designs are required where each participant obtains multiple observations of these interactions, which would require replication at the target level of inference (i.e., each participant completes the entire two-phase intervention twice). Thus, there may be unknown bias in our variance estimates and target inferences thereof if the assumption of consistency does not hold.

However, as both conditions are completed concurrently at the participant-level, these constraints of typical two-phase crossover designs may be of less relevance.

To validate our assumptions and the collective set of novel methods used for inference, we created an R shiny application which performs simulations with these techniques to ensure their ability to detect IRV when it is known to be present or not. Further details (i.e., link to the application and code) can be accessed in the supplementary materials (https://osf.io/d5cxy/).

### Statistical Analyses

#### Test-Retest Reliability

Two-way-random intraclass correlation coefficients (ICC), standard errors of measurement (SEM), and coefficients of variation (CV) were calculated for each outcome using the SimplyAgree package (Caldwell, 2022) to assess the test-retest reliability of each outcome. For CSA, MT, and MVIC the two measurements performed at the same occasion were compared. For 1RM the differences between limbs at the baseline measurement of both phases were compared; which assumes a negligible systematic difference between limbs. These estimates help to roughly approximate the degree to which phase-to-phase variability in the training effects are due to measurement error versus sampling variance and biological variability.

#### Gross Variability

The gross variability of each outcome was quantified via independently calculated change scores for each limb in each phase with missing values imputed with the cluster-specific mean (i.e., limb nested within participant). For gross variability relevant to GEN IRV the change scores of a participant’s limbs within a given phase were averaged. Alternatively, for gross variability relevant to CON IRV, the change scores remained on the limb-level to maintain the integrity of the protocols they were assigned to. Waterfall, line, and diagonal plots were then produced for each outcome which compare the results to a threshold derived from the estimate of measurement error (SEM x 2). These plots visualize the inter-individual variability without accounting for confounding sources of variation (i.e., sampling variance, biological variability, etc.) and allow for comparison to previous work (Hubal *et al*., 2005; Erskine *et al*., 2010; Ahtiainen *et al*., 2016; Damas *et al*., 2019; Hammarström *et al*., 2020; Lixandrão *et al*., 2024).

#### Naïve Estimation of Inter-individual Response Variation

##### Magnitude

Using the limb-specific change scores from the examination of gross variability, individual training effects (ITE) were calculated for each GEN and CON, respectively. For GEN, the ITE were estimated by first calculating the average of the limb-specific change scores within each phase (i.e., phase-specific training effect) such that there were two values per participant. These values were then averaged across phases to derive the ITE (i.e., one value per participant). For CON, the ITE were estimated by first calculating the difference between HV and LV limb-specific change scores in each phase (i.e., phase-specific training effect) such that there were two values per participant. These values were then averaged across phases to derive the ITE (i.e., one value per participant).

To calculate the average training effect (ATE), all of the ITE were averaged for each GEN and CON, respectively. Further, to mirror the structure of the statistical model used for integrated estimation (which included interaction terms to proxy potential carryover effects), phase-specific means were calculated by averaging the phase-specific training effects for each GEN and CON, respectively. Then, the ATE was subtracted from each of these phase specific means to obtain the aforementioned correction for potential carryover effects. These corrections were used to better estimate the remaining WPV by calculating the standard deviation of the differences between phase-specific means and the ITE after the phase-specific correction was applied (i.e., residuals).

The average WPV was then utilized to construct 95% confidence intervals (CI) around each ITE. A common variance was chosen for the individual-level estimates as participant-specific variances are particularly unstable in small samples (Senn, 2024), though this choice does assume true homogeneity. Further, for the ATE, the standard deviation of the ITE was divided by the square root of the sample size to obtain the standard error. Standard errors for prediction intervals (PI) were obtained for the ATE by first summing the squares of the standard error and the average WPV, and then taking the square root. Then 95% CI were obtained by multiplying the previously specified standard errors by the t-value associated with the given degrees of freedom.

Once the initial calculations had been performed the magnitude of the standard deviations associated with IRV and WPV could be estimated (i.e., SD_IRV_ and SD_WPV_). Specifically, the standard deviation of the ITE was considered the naïve estimate of SD_IRV_, while the average WPV was considered the naïve estimate of SD_WPV_. Confidence intervals were constructed for both SD_IRV_ and SD_WPV_ via nonparametric bootstrapping with 1000 replicates.

##### Reliability

Using the phase-specific training effects from each participant, reliability was established. Specifically, a Pearson’s correlation coefficient (r) was calculated by estimating the slope between the standardized training effects of phases 1 and 2. The relationship was visualized by fitting a linear regression model on the unstandardized training effects of phases 1 and 2 and extracting the predictions thereof (i.e., means and 95% CI). Prediction intervals (PI) were constructed by integrating the residual error term of the model into the standard error of the mean (i.e., √SE² + σ²) and then multiplying by the t-value associated with the corresponding degrees of freedom. Pearson’s r helps to establish if a given domain of IRV is “detectable” or “real”. A strong correlation would suggest that the ITE are “reliable” across phases, with the precision of the correlation roughly indicating the magnitude of SD_WPV_. Alternatively, a weak correlation would suggest that the ITE are not “reliable” across phases. Additionally, we constructed intraclass correlation coefficients (ICC) that estimate the proportion of total variance explained by IRV. Specifically the square of SD_IRV_ was divided by the total variance (i.e., SD_IRV_^2^ + SD_WPV_^2^) to calculate the ICC. The ICC roughly approximates the signal-to-noise ratio of the variances, with higher proportions suggesting a stronger signal of IRV. Confidence intervals were constructed for both Pearson’s r and the ICC via nonparametric bootstrapping with 1000 replicates.

##### Agreement

To contextualize if a given estimate of IRV is “meaningful” or “actionable”, we assessed the agreement of the training effects. We constructed modified versions of Bland Altman plots to visualize the competing sources of variation (i.e., SD_IRV_ and SD_WPV_). Specifically, we plotted the difference between the ITE and ATE (i.e., x-axis) against the difference between the phase-specific means from the ITE after the phase-specific correction was applied to account for potential carryover effects (i.e., y-axis). Limits of agreement (LOA) were calculated by multiplying the SD_WPV_ by the t-value associated with the corresponding degrees of freedom. As these values represent deviation from the mean, the “bias” typically estimated in agreement analysis is necessarily zero, but the relative spread of data points (i.e., height versus width) is of paramount interest. The relative dispersion of data identifies the predominant source of variation in the ITE. Data that is predominantly spread horizontally signals that the differences between ITE (i.e., SD_IRV_) exceeds the variability that occurs from phase to phase (i.e., SD_WPV_), indicating meaningful or actionable IRV. Alternatively, data that is predominantly spread vertically signals that the differences between the ITE is negligible (i.e., SD_IRV_) compared to the remaining WPV (i.e., SD_WPV_), indicating IRV unlikely to be meaningful or actionable. To formally quantify the differences between these variance components, both multiplicative and additive contrasts were performed. Contrasts on both scales (i.e., multiplicative and additive) were performed to detect potential sensitivity in the comparison utilized (Nakagawa *et al*., 2014; Mills *et al*., 2021). Multiplicative log variability ratios (lnVR) were calculated as the ratio between SD_IRV_ and SD_WPV_ with values greater than zero indicating that IRV exceeds the remaining WPV. Similarly, additive differences in variability (DiV) were calculated as the difference between the squares of SD_IRV_ and SD_WPV_. Again, values greater than zero indicate that IRV exceeds the remaining WPV. Confidence intervals were constructed for LOA, lnVR, and DiV via nonparametric bootstrapping with 1000 replicates.

#### Integrated Estimation of Inter-individual Response Variation

##### Magnitude

To formally estimate the magnitude of effects relevant to the primary research questions, we fit Bayesian multivariate mixed effect linear regression models using the *brms* package (Bürkner, 2017). These models were parameterized as an extension of an analysis of covariance (ANCOVA) with an adjustment for the baseline value of the dependent variable. Fitting the model in this way allowed for multiple outcomes, all three timepoints in each phase, and multiple observations per time point to be modeled collectively which improves statistical power/precision and allow for better estimates of the target variance components.

Hypertrophy (i.e., CSA and MT) and strength (i.e., 1RM and MVIC) outcomes were modeled separately, but the outcomes within each model were estimated jointly and assumed to have a correlation between their variances (i.e., residual errors and random effects). Further, each model included fixed effects for time, phase, sex, and time-condition, time-phase, time-condition-phase interactions. This fixed effect structure allows the model to estimate the target estimands (i.e., main effect of time and time-condition interaction) as well as adjust for potential bias due to an ineffective washout period (i.e., main effect of phase) and differences in the training effect between phases which may proxy carryover effects (i.e., time-phase and time-condition-phase interactions).

To account for the dependent observations, nested random intercepts were included for each participant and their lower limbs. To determine the optimal random slope structure, multiple multivariate models were initially fit and compared using the *lme4* package for ease of computation (Bates *et al*., 2015). The first candidate model included a maximal random slope structure (i.e., all within-participant factors and their consequent interactions) while the other models gradually reduced the number of random slopes included. This approach was chosen as it seems to offer a better balance of type 1 and type 2 error compared to an underparameterized version of the model for a given data generating process (Barr *et al*., 2013; Matuschek *et al*., 2017; Oberauer, 2022). However, after comparing model performance with the *bayestestR* package (i.e., Bayesian information criteria approximated Bayes factors) (Makowski *et al*., 2019), all random slopes aside from those necessary to estimate the target variance components (i.e., random slopes of time and time-condition interaction on the participant level) were removed in favor of model parsimony. For both models random effects were assumed to follow normal distributions with a mean of zero and an unknown variance to be estimated, while residual errors (i.e., WPV) were assumed to be homogeneous across clusters (i.e., participants and limbs nested within participants) and all strata of fixed effects. These are standard statistical assumptions but are of particular relevance to the primary research questions. Due to the novelty of the study design and the exploratory nature of the analysis, each model was fit with the default weakly regularizing priors. Four Monte Carlo Markov Chains with 1000 warmup and 4000 sampling iterations were used for each model. The quality of each model was deemed sufficient upon inspection of the density, trace, and residual plots created from draws of the posterior distribution.

Adjusted training effects for each outcome were calculated using the *marginaleffects* (Arel-Bundock, 2023) and *tidybayes* packages (Kay, 2023). Specifically, estimates were generated by extracting draws from the posterior predictive distributions (i.e., includes observation-level error) and the expected values thereof (i.e., removes observation-level error). For group-level estimates, all random effects were set to zero to reflect the grand mean; however, for individual-level estimates, all random effects were included to preserve the variances estimated at each level in the model. The draws of each estimate were then summarized with point estimates of central tendency (i.e., mode), the 95% highest density intervals of the posterior predictions (HDPI) and expected values thereof (HDI). Finally, for each estimate, the posterior probability (p) of the effect being in the direction consistent with the observed estimate was calculated (i.e., P(θ > 0) for positive estimates and P(θ < 0) for negative estimates).

For GEN, training effects were calculated as the change in the outcome proportionally marginalized across sex, phase, and condition (i.e., main effect of time). Evidence of potential carryover effects were proxied by the difference in training effects between phases (i.e., time-phase interaction), proportionally marginalized across sex and condition. For CON, training effects were calculated as the difference in the changes in the outcome between conditions proportionally marginalized across sex and phase (i.e., time-condition interaction). Evidence of potential carryover effects were proxied by the difference in training effects between phases (i.e., time-condition-phase interaction), proportionally marginalized across sex.

In order to derive valid estimates of SD_IRV_ and SD_WPV_ for each domain (i.e., GEN and CON), respectively; additional steps are required. Because conditional estimates (i.e., estimates from the existing sample of participants) tend to underestimate the true variance in the implied population, training effects with and without observation level error were calculated for 100 participants simulated from the models parameters. The standard deviation of these simulated training effects without observation level error represents the unbiased estimate for SD_IRV_. Alternatively, the standard deviation of the differences between the simulated training effects with and without observation level error (i.e., residuals) represents the unbiased estimate for SD_WPV_.

##### Reliability

Using the posterior draws of the phase-specific training effects from each of the simulated participants, reliability was established. Specifically, Pearson’s r was calculated by estimating the slope between the standardized training effects of phases 1 and 2 within in draw. The relationship was visualized by fitting a linear regression model on the unstandardized training effects of phases 1 and 2 within each posterior draw and extracting the predictions thereof. Predictions with observation-level error were generated by extracting the residual error term from each model and simulating offsets to add to the predictions using a normal distribution with a mean of 0 and variance in line with the estimate of the error term. Then both credible and prediction intervals were created by summarizing the posterior draws for the estimates with and without observation-level error. Similarly, we calculated the ICC in each posterior draw as the square of SD_IRV_ was divided by the total variance (i.e., SD_IRV_^2^ + SD_WPV_^2^), consequently summarizing those draws.

##### Agreement

To contextualize if a given estimate of IRV is “meaningful” or “actionable”, we assessed the agreement of the training effects using the posterior predictive draws and their expected values from each of the simulated participants. We constructed modified versions of Bland Altman plots to visualize the competing sources of variation (i.e., SD_IRV_ versus SD_WPV_). Specifically, we plotted the difference between the expected values of the ITE and ATE (i.e., x-axis) against the difference between the phase-specific training calculated from the posterior predictive distribution (i.e., with observation-level error) and the expected value of the ITE in a given phase (i.e., y-axis). Uncertainty intervals were constructed by summarizing the 95% HDI’s for the expected values of the differences (i.e., interval for bias) and the posterior predictions with observation-level error (i.e., limits of agreement). To formally quantify the differences between these variance components, both multiplicative and additive contrasts were performed. Using the posterior draws of SD_IRV_ and SD_WPV_, lnVR were calculated with values greater than zero indicating that IRV exceeds the remaining WPV. Similarly, additive DiV’s were calculated using the posterior draws as the difference between the squared values of SD_IRV_ and SD_WPV_.

#### Prediction

If sufficient evidence of IRV was identified, we planned to explore potential candidate predictors of this variability in training effects. Specifically, we planned to use the posterior draws of the expected values of the differences between the ITE and ATE as this directly represents the variability explained by IRV. Within each draw, we planned to fit linear regression models with the differences and baseline predictor of interest. From there we would calculate the slope and then summarize these estimates across the posterior draws to determine the ability of the predictor to directionally identify IRV. However, since our primary analyses did not reveal indisputable evidence of meaningful IRV, we did not proceed with these predictive models, as there would be insufficient variability to explain. Further, even if potential candidate predictors were identified, further research would need to be dedicated to evaluating the true causal effect of these variables by allocating individuals in concordance with the predictor(s) and verifying the intended impact (e.g., magnitude, precision, out of sample predictive power).

#### Simulation

To validate our analytical framework, we created an R shiny application that performs simulations to assess the ability of our methods to detect IRV when known to be absent, present, or actionable (https://osf.io/d5cxy/). The application simulates data with customizable parameters for both traditional crossover and replicated crossover designs. Then gross, naive, and integrated (i.e., adjusted) estimates were calculated for each design. For each approach, we quantified the ITE, SD_IRV_, SD_WPV_, Pearson’s r, ICC, LOA, lnVR, and DiV. These metrics can be compared against their true population parameters to assess bias and precision. We also explored the ability of model comparisons via Bayesian information criteria to approximate Bayes factors (Log BF) to identify whether IRV was present in the data generating process or not, probabilistically. To conceptually demonstrate the ability of our methodology to recover known parameters, we have included an example in the application where the data was simulated such that IRV was know to be absent while maintaining some WPV (i.e., SD_WPV_ > SD_IRV_ = 0). Alternatively, one could perform simulations where true IRV is present but not actionable (i.e,. SD_WPV_ > SD_IRV_ > 0), or where true IRV is both present and actionable (i.e,. SD_WPV_ < SD_IRV_ > 0) by slightly altering the simulation parameters.

## RESULTS

### Hypertrophy

#### Test-Retest Reliability

The ICC for VL CSA was 0.98 [95% CI: 0.97, 0.99] with a SEM of 1.79 cm^2^ [95% CI: 1.52, 2.17] and CV of 2.77% [95% CI: 2.36, 3.35]. For VL MT, the ICC was 0.97 [95% CI: 0.96, 0.98] with a SEM of 0.56 mm [95% CI: 0.48, 0.68] and CV of 2.12% [95% CI: 1.81, 2.57]. These results can be visualized in Figure 2.

**Figure 2:**
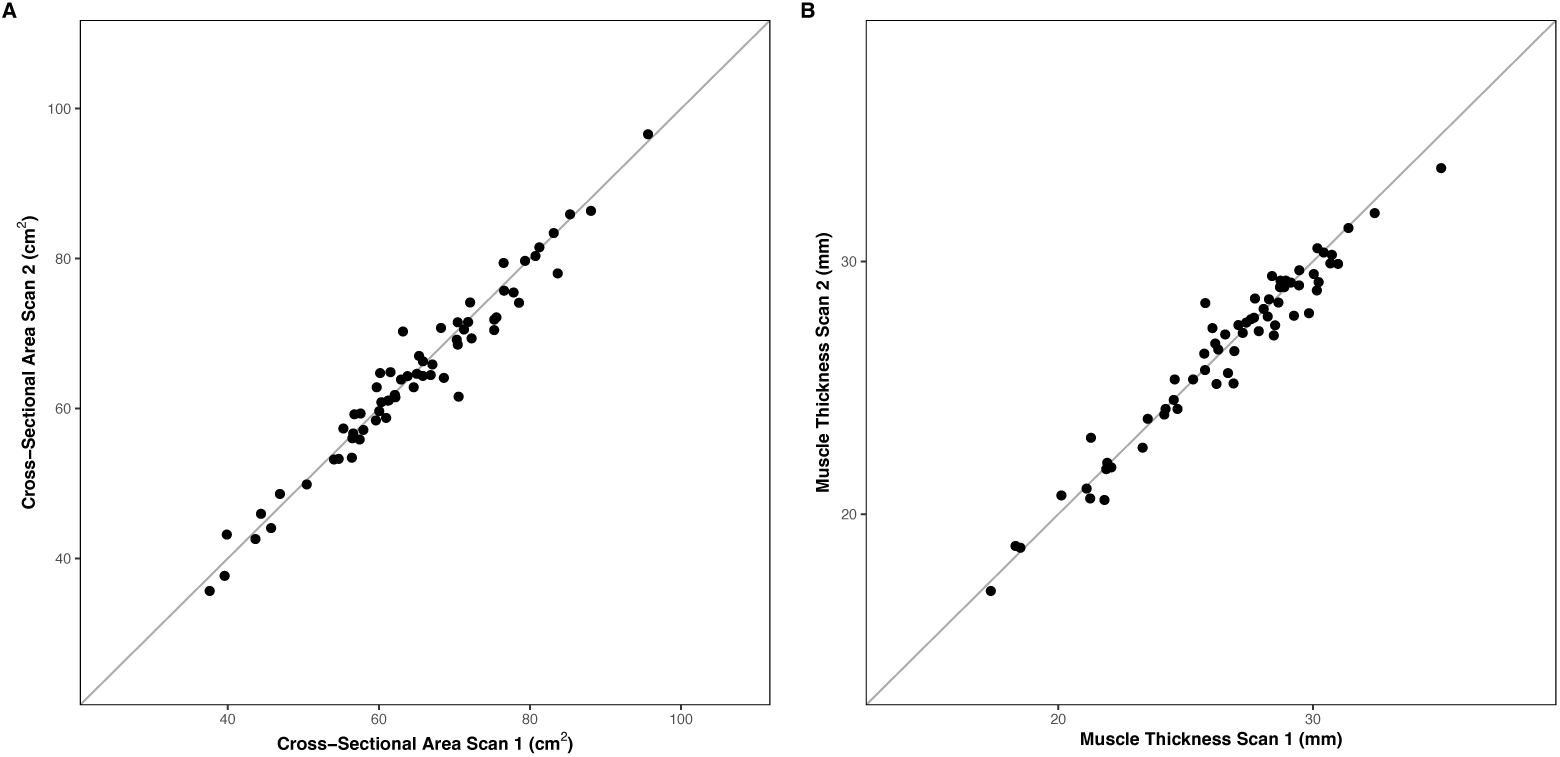
Test-retest reliability of vastus lateralis (VL) A) cross-sectional area (CSA) and B) muscle thickness (MT). The x-axis represents the value obtained at the first measurement occasion, while the y-axis represents the value obtained at the second occasion.

#### Gross Variability

For GEN, there was a range of -3.65 cm^2^ to 17.38 cm^2^. When treating each phase independently, 29 participants demonstrated positive changes in VL CSA while 3 participants experienced negative change scores in VL CSA.

For CON, there was a range of -1.44 cm^2^ to 20.89 cm^2^ for HV conditions and -10.69 cm^2^ to 18.8 cm^2^ for LV conditions. When treating each phase independently, 22 participants demonstrated superior changes in the HV condition while 10 participants experienced superior change scores with the LV condition. These results are visualized in Figure 3.

**Figure 3:**
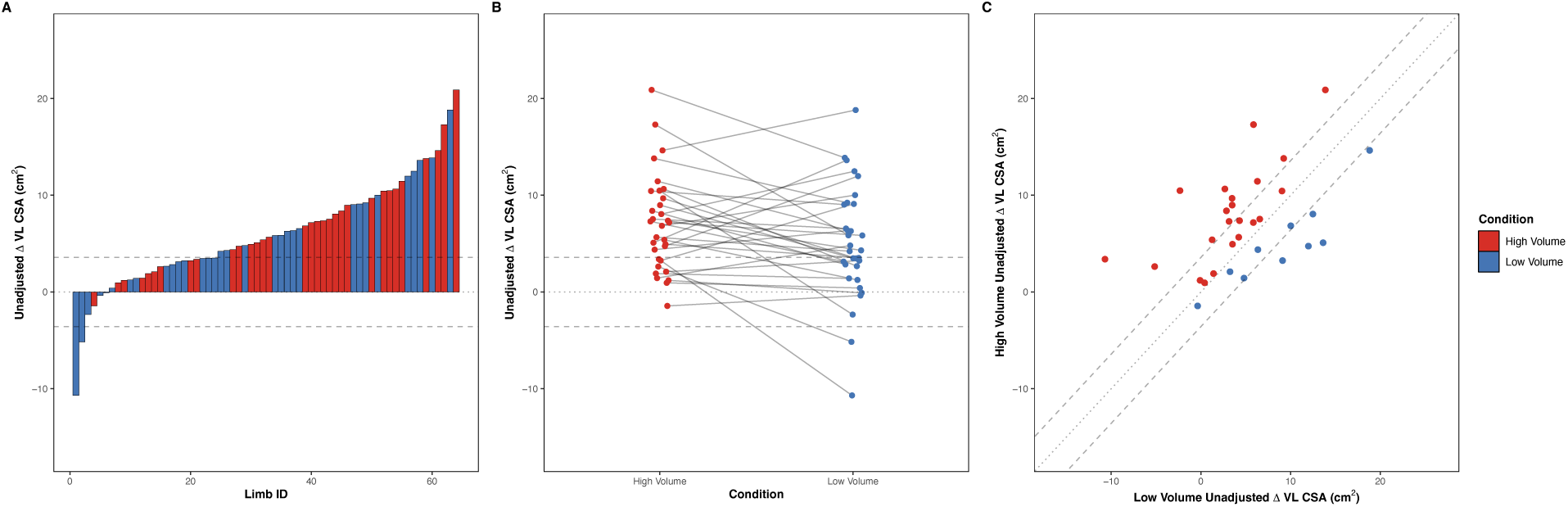
Gross variability of unadjusted change scores in vastus lateralis cross-sectional area (VL CSA) following resistance training with high (red) or low (blue) weekly set volumes. The gray dashed lines represent the threshold derived from the estimate of measurement error (standard error of measurement (SEM) x 2) relative to a null change. A) waterfall plot in which each bar represents the unadjusted change score of a given limb, B) line plot in which the unadjusted change scores of each participant within a given phase is connected by a gray line, c) diagonal plot in which the x-axis represents the unadjusted change score of the low volume condition while the y-axis represents the high volume condition for a given participant. The dark diagonal line is the line of identity which represents a perfect correlation in the change scores between conditions. Red points are those who favor high volumes and blue points are those who favor low volumes.

The results for VL MT largely mirrored the primary outcome. Specifically for GEN, there was a range of -0.93 mm to 5.26 mm. When treating each phase independently, 29 participants demonstrated positive changes in VL MT while 3 participants experienced negative change scores in VL MT.

For CON, there was a range of -2.41 mm to 6.37 mm for HV conditions and -2.12 mm to 5.7 mm for LV conditions. When treating each phase independently, 19 participants demonstrated superior changes in the HV condition while 13 participants experienced superior change scores with the LV condition. These results can be visualized in the supplementary materials.

#### Naïve Estimates of Inter-individual Response Variation

##### Magnitude

For GEN, the naïve estimate of the ATE for VL CSA was 6.03 cm^2^ [95% CI: 3.9, 8.15; 95% PI: - 1.76, 13.81]. On the participant level, ITE ranged from -1.55 cm^2^ to 13.82 cm^2^ with the 95% CI of 13 out of 16 individuals suggesting estimates were compatible with the ATE. These results can be visualized in Figure 4A. The resulting estimate of SD_IRV_ was 3.99 cm^2^ [95% CI: 2.55, 5.43] while SD_WPV_ was 3.53 cm^2^ [95% CI: 2.39, 4.67].

**Figure 4:**
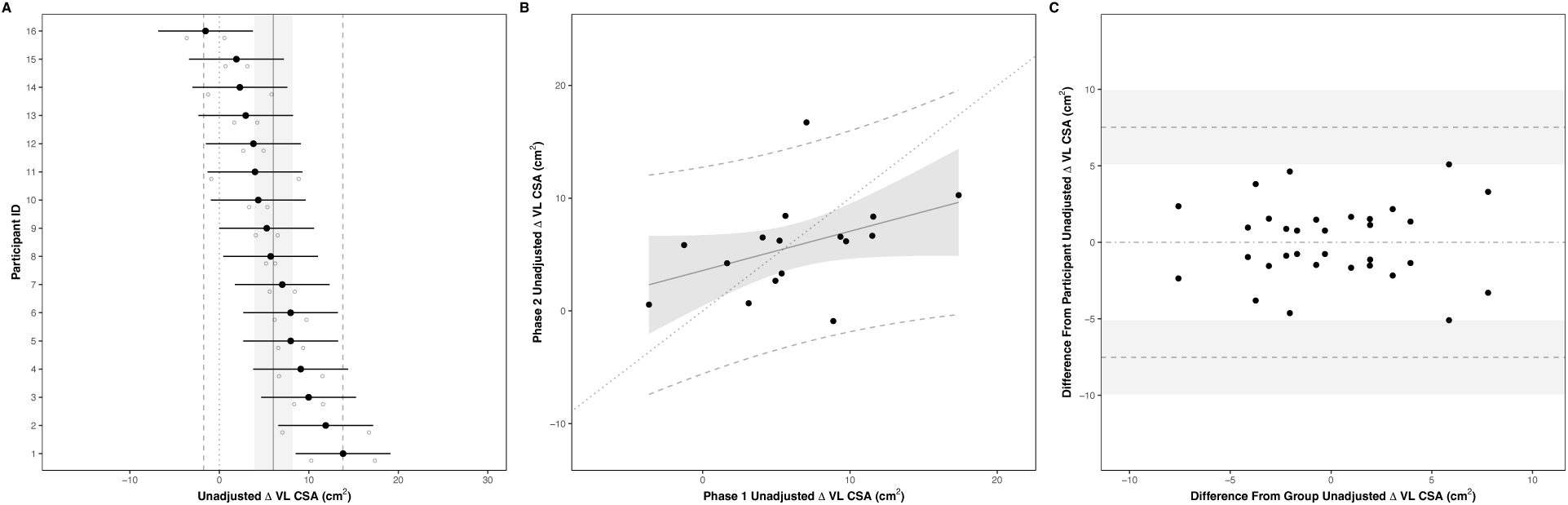
Naïve estimates of general (GEN) individual response variation (IRV) in vastus lateralis cross-sectional area (VL CSA) following resistance training. The gray line shaded region and dashed lines represent the average training effect (ATE), 95% confidence, and prediction intervals, respectively. A) Forest plot of individual training effects (ITE). Black dots represent a given participants’ mean and uncertainty interval, while open circles represent the observed change score from each phase. B) Reliability of training effects between phases. Black dots represent the intersection of observed change scores from each phase. C) Agreement of training effects between phases. X-axis represents the deviation of ITE from the ATE, while the y-axis represents the deviation of the observed change score from each phase from their respective ITE.

For CON, the naïvely estimated ATE of VL CSA was 2.11 cm^2^ [95% CI: 0.39, 3.82; 95% PI: -10.32, 14.53]. On the participant level, ITE ranged from -2.23 cm^2^ to 7.69 cm^2^ with the 95% CI of 16 out of 16 individuals suggesting estimates were compatible with the ATE. These results can be visualized in Figure 5A. The resulting estimate of SD_IRV_ was 3.21 cm^2^ [95% CI: 2.34, 4.09] while SD_WPV_ was 6.33 cm^2^ [95% CI: 4.39, 8.27].

**Figure 5:**
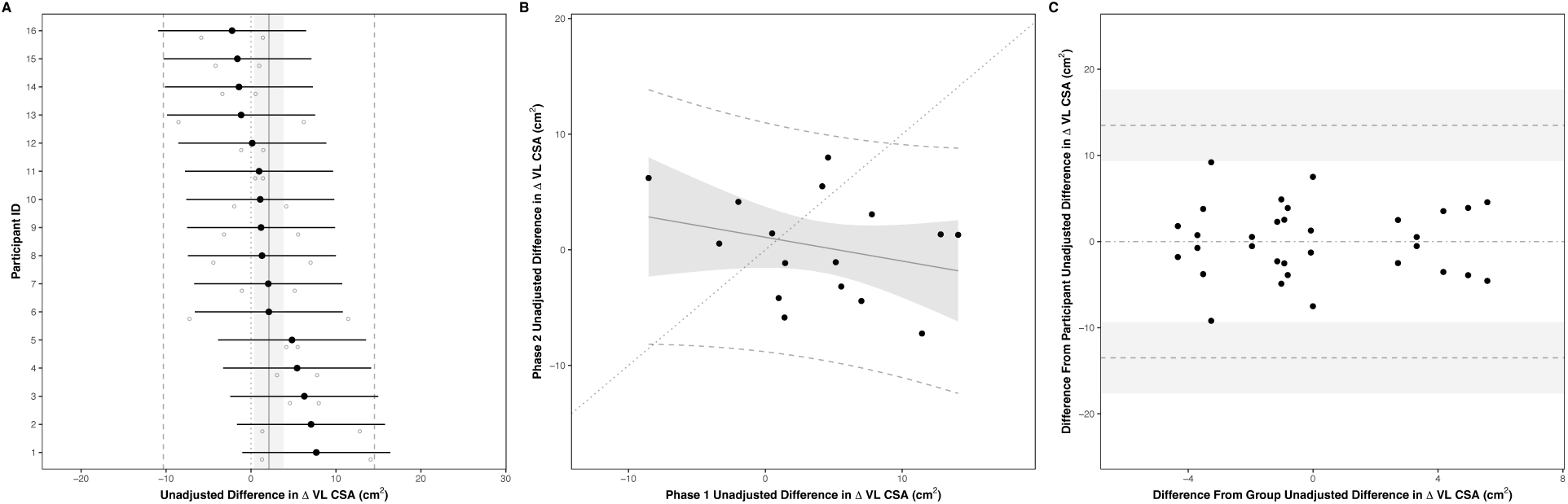
Naïve estimates of condition-specific (CON) individual response variation (IRV) in vastus lateralis cross-sectional area (VL CSA) following resistance training. The gray line shaded region and dashed lines represent the average training effect (ATE), 95% confidence, and prediction intervals, respectively. A) Forest plot of individual training effects (ITE). Black dots represent a given participants’ mean and uncertainty interval, while open circles represent the observed change score from each phase. B) Reliability of training effects between phases. Black dots represent the intersection of observed change scores from each phase. C) Agreement of training effects between phases. X-axis represents the deviation of ITE from the ATE, while the y-axis represents the deviation of the observed change score from each phase from their respective ITE.

The results for VL MT largely mirrored the primary outcome. For GEN, the naïvely estimated ATE was 1.66 mm [95% CI: 1.06, 2.26; 95% PI: -1.14, 4.46]. On the participant level, ITE ranged from -0.23 mm to 4.08 mm with the 95% CI of 15 out of 16 individuals suggesting estimates were compatible with the ATE. The resulting estimate of SD_IRV_ was 1.12 mm [95% CI: 0.73, 1.51] while SD_WPV_ was 1.29 mm [95% CI: 0.63, 1.95].

For CON, the naïvely estimated ATE of VL MT was 0.58 mm [95% CI: -0.11, 1.28; 95% PI: -4.79, 5.96] indicating an advantage to the HV condition on the group-level. On the participant level, ITE ranged from -3.08 mm to 2.61 mm with the 95% CI of 16 out of 16 individuals suggesting estimates were compatible with the ATE. The resulting estimate of SD_IRV_ was 1.3 mm [95% CI: 0.56, 2.03] while SD_WPV_ was 2.6 mm [95% CI: 1.83, 3.37]. These results can be visualized in the supplementary materials.

##### Reliability

For GEN, the average difference between phases in the ATE of VL CSA was -0.51 cm^2^ [95% CI: - 3.25, 2.22; 95% PI: -8.48, 7.45]. On the participant level, phase-specific training effects ranged from -1.29 cm^2^ to 14.08 cm^2^ in phase 1 and -1.81 cm^2^ to 13.56 cm^2^ in phase 2. These results can be visualized in Figure 4B. The resulting estimate of Pearson’s correlation coefficient was 0.42 [95% CI: 0.03, 0.82] while the ICC was 0.56 [95% CI: 0.37, 0.75].

For CON, the average difference between phases in the ATE of VL CSA was -3.66 cm^2^ [95% CI: - 8.16, 0.83; 95% PI: -16.77, 9.44]. On the participant level, phase-specific training effects ranged from -8.53 cm^2^ to 14.08 cm^2^ in phase 1 and -7.24 cm^2^ to 7.98 cm^2^ in phase 2. These results can be visualized in Figure 5B. The resulting estimate of Pearson’s correlation coefficient was -0.28 [95% CI: -0.75, 0.19] while the ICC was 0.2 [95% CI: 0.07, 0.34].

The results for VL MT largely mirrored the primary outcome. For GEN, the average difference between phases in the ATE of VL MT was 0.14 mm [95% CI: -0.86, 1.14; 95% PI: -2.77, 3.06]. On the participant level, phase-specific training effects ranged from -0.31 mm to 4 mm in phase 1 and -0.16 mm to 4.15 mm in phase 2. The resulting estimate of Pearson’s correlation coefficient was 0.18 [95% CI: -0.52, 0.87] while the ICC was 0.43 [95% CI: 0.12, 0.74].

For CON, the average difference between phases in the ATE of VL MT was -0.99 mm [95% CI: - 2.94, 0.96; 95% PI: -6.66, 4.68]. On the participant level, phase-specific training effects ranged from -3.8 mm to 4.35 mm in phase 1 and -3.26 mm to 3.26 mm in phase 2. The resulting estimate of Pearson’s correlation coefficient was -0.33 [95% CI: -0.93, 0.26] while the ICC was 0.2 [95% CI: -0.02, 0.42]. These results can be visualized in the supplementary materials.

##### Agreement

For GEN, the differences between the ITE and the ATE for VL CSA ranged from -5.09 cm^2^ to 5.09 cm^2^. Alternatively, the difference between the ITE with and without observation-level error ranged from -5.09 cm^2^ to 5.09 cm^2^. Consequently the limits of agreement extended from 7.53 cm^2^ [95% CI: 5.1, 9.95] to -7.53 cm^2^ [95% CI: -5.1, -9.95]. These results can be visualized in Figure 4C. The resulting estimate of lnVR was 0.25 [95% CI: -0.54, 1.03], while DiV was 3.47 [95% CI: -7.27, 14.2].

For CON, the differences between the ITE and the ATE for VL CSA ranged from -9.2 cm^2^ to 9.2 cm^2^. Alternatively, the difference between the ITE with and without observation-level error ranged from -9.2 cm^2^ to 9.2 cm^2^. Consequently the limits of agreement extended from 13.49 cm^2^ [95% CI: 9.36, 17.63] to -13.49 cm^2^ [95% CI: -9.36, -17.63]. These results can be visualized in Figure 5C. The resulting estimate of lnVR was -1.36 [95% CI: -2.22, -0.49], while DiV was -29.74 [95% CI: -54.17, -5.3].

The results for VL MT largely mirrored the primary outcome. For GEN, the differences between the ITE and the ATE for VL MT ranged from -2.71 mm to 2.71 mm. Alternatively, the difference between the ITE with and without observation-level error ranged from -2.71 mm to 2.71 mm. Consequently the limits of agreement extended from 2.74 mm [95% CI: 1.34, 4.15] to -2.74 mm [95% CI: -1.34, -4.15]. The resulting estimate of lnVR was -0.28 [95% CI: -1.65, 1.08], while DiV was -0.41 [95% CI: -2.35, 1.54].

For CON, the differences between the ITE and the ATE for VL MT ranged from -3.01 mm to 3.01 mm. Alternatively, the difference between the ITE with and without observation-level error ranged from -3.01 mm to 3.01 mm. Consequently the limits of agreement extended from 5.53 mm [95% CI: 3.89, 7.17] to -5.53 mm [95% CI: -3.89, -7.17]. The resulting estimate of lnVR was -1.39 [95% CI: -2.96, 0.19], while DiV was -5.05 [95% CI: -9.69, -0.41]. These results can be visualized in the supplementary materials.

#### Integrated Estimates of Inter-individual Response Variation

##### Magnitude

For GEN, the integrated estimate of ATE of VL CSA was 6.01 cm^2^ [95% HDI: 4.11, 8.07; 95% HDPI: 1.11, 11.09] with a 100% posterior probability of exceeding the null. On the participant level, ITE ranged from 1.34 cm^2^ to 11.95 cm^2^ with the 95% HDI of 10 out of 16 individuals suggesting estimates were compatible with the ATE. These results can be visualized in Figure 6A and 7A, respectively. The resulting estimate of SD_IRV_ was 3.43 cm^2^ [95% HDI: 2.06, 5.39] with a 100% posterior probability of exceeding the null. The estimate of SD_WPV_ was 1.63 cm^2^ [95% HDI: 1.38, 1.91] with a 100% posterior probability of exceeding the null.

**Figure 6:**
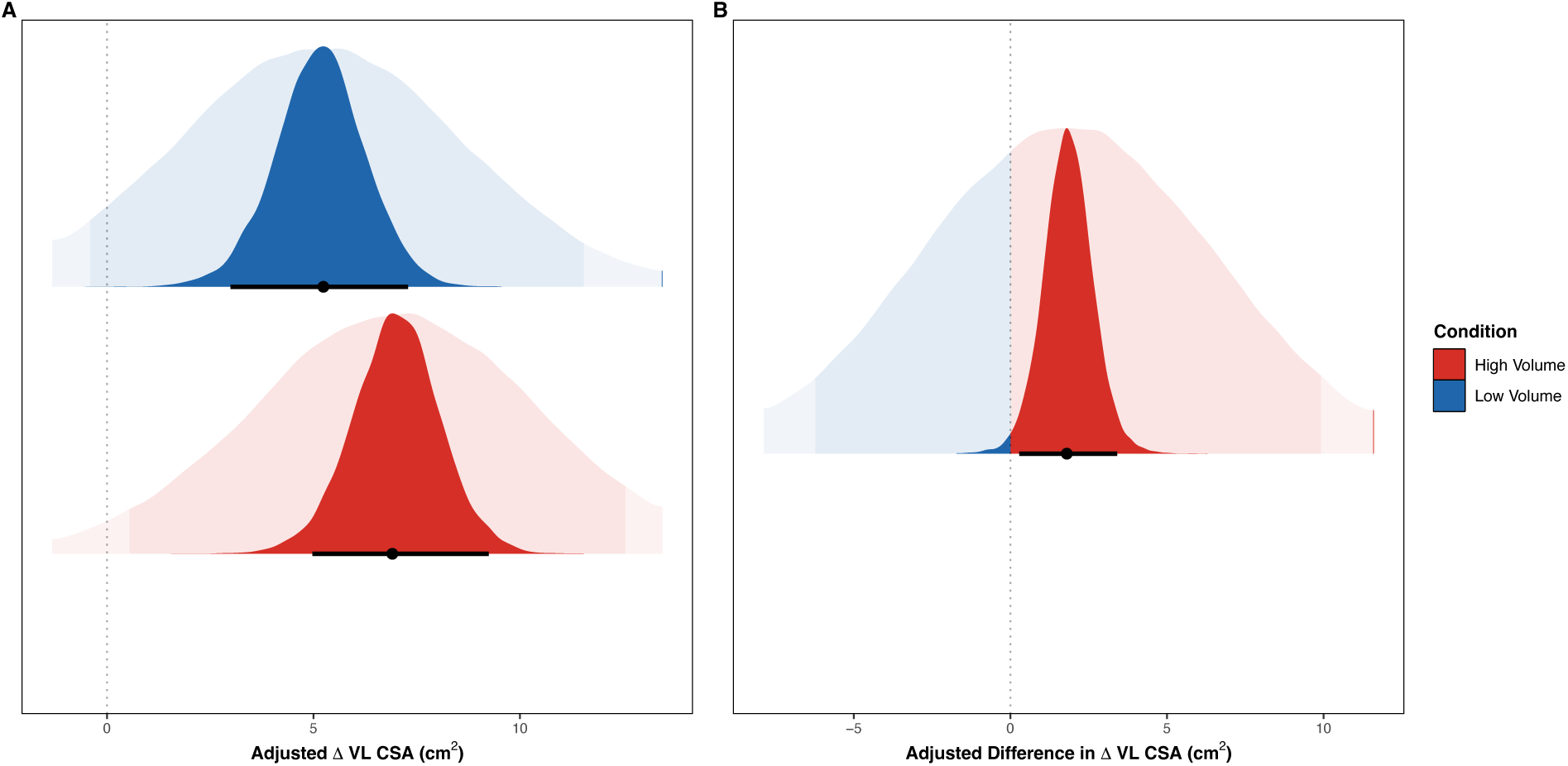
Adjusted average training effects (ATE) for changes in vastus lateralis cross-sectional area (VL CSA) A) within, and B) between conditions. Black dots and intervals represent the mode and the 95% highest density intervals (HDI) of the posterior distribution while the faded densities in the background represent the draws from the posterior predictive distribution, where the darker region represents the 95% prediction interval. For within-condition effects, the red distributions represents high volumes (HV) and the blue distributions represents low volumes (LV). For the between-condition effect, the red portion of the distributions represents the posterior draws that favor HV while the blue portion of the distributions favor LV.

**Figure 7:**
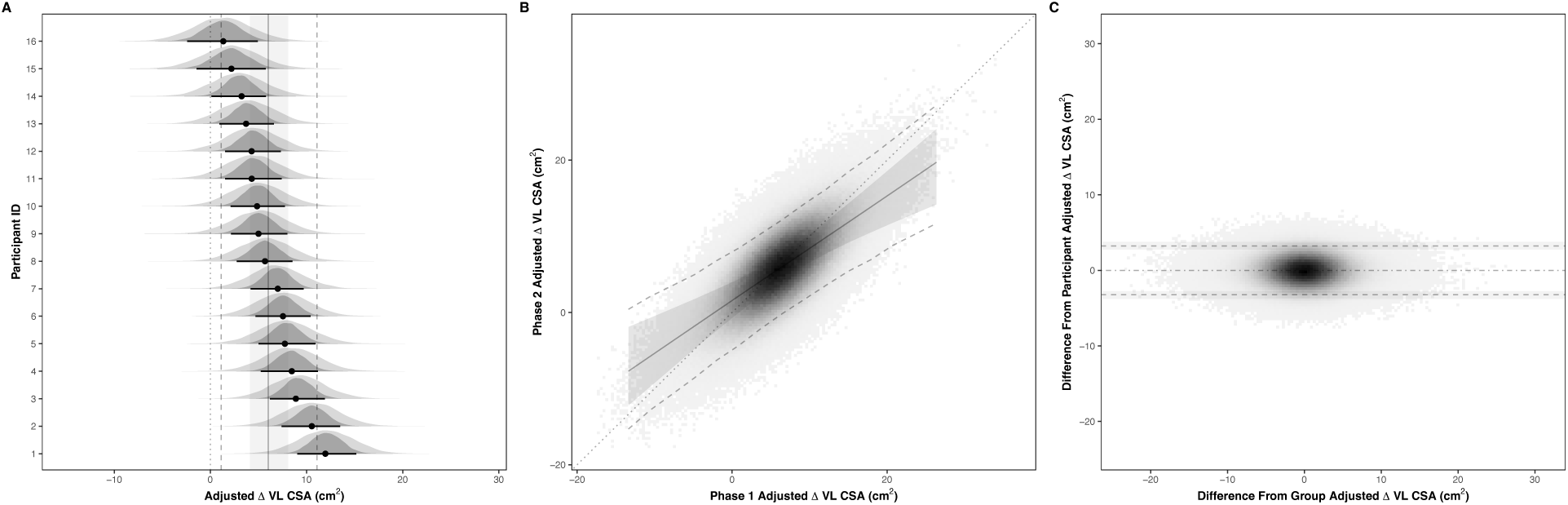
Integrated estimates of general (GEN) of individual response variation (IRV) in vastus lateralis cross-sectional area (VL CSA) following resistance training. The gray line shaded region and dashed lines represent the average training effect (ATE), 95% highest density, and prediction intervals, respectively. A) Forest plot of the posterior distributions of individual training effects (ITE). Black dots and uncertainty intervals represent the participants’ mode and uncertainty interval while the faded densities represent the posterior predictive distribution. B) Reliability of training effects between phases. Colored bins represent the intersection posterior predictions of training effects of each phase from simulated participants. C) Agreement of training effects between phases. X-axis represents the deviation of simulated ITE from the ATE, while the y-axis represents the deviation of the simulated posterior predictions from each phase from their respective ITE.

For CON, the integrated estimate of ATE of VL CSA was 1.8 cm^2^ [95% HDI: 0.29, 3.41; 95% HDPI: -7.41, 11.17] with a 98.77% posterior probability of exceeding the null. On the participant level, ITE ranged from 1.58 cm^2^ to 2.08 cm^2^ with the 95% HDI of 16 out of 16 individuals suggesting estimates were compatible with the ATE. These results can be visualized in Figures 6B and 8A, respectively. The resulting estimate of SD_IRV_ was 0.37 cm^2^ [95% HDI: 0, 3.52] with a 100% posterior probability of exceeding the null. The estimate of SD_WPV_ was 3.28 cm^2^ [95% HDI: 2.78, 3.82] with a 100% posterior probability of exceeding the null.

**Figure 8:**
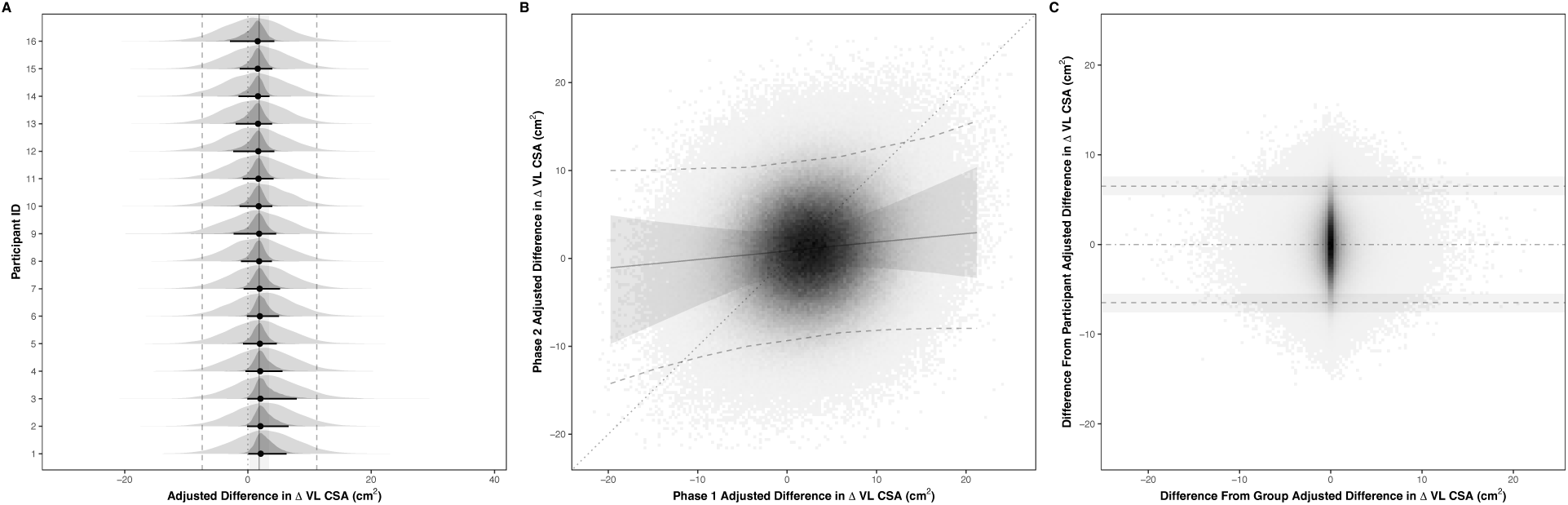
Integrated estimates of condition-specific (CON) individual response variation (IRV) in vastus lateralis cross-sectional area (VL CSA) following resistance training. The gray line shaded region and dashed lines represent the average training effect (ATE), 95% highest density, and prediction intervals, respectively. A) Forest plot of the posterior distributions of individual training effects (ITE). Black dots and uncertainty intervals represent the participants’ mode and uncertainty interval while the faded densities represent the posterior predictive distribution. B) Reliability of training effects between phases. Colored bins represent the intersection posterior predictions of training effects of each phase from simulated participants. C) Agreement of training effects between phases. X-axis represents the deviation of simulated ITE from the ATE, while the y-axis represents the deviation of the simulated posterior predictions from each phase from their respective ITE.

The results for VL MT largely mirrored the primary outcome. For GEN, the integrated estimate of ATE of VL MT was 1.73 mm [95% HDI: 1.19, 2.22; 95% HDPI: 0.24, 3.18] with a 100% posterior probability of exceeding the null. On the participant level, ITE ranged from 0.58 mm to 3.44 mm with the 95% HDI of 12 out of 16 individuals suggesting estimates were compatible with the ATE. The resulting estimate of SD_IRV_ was 0.84 mm [95% HDI: 0.5, 1.4] with a 100% posterior probability of exceeding the null. The estimate of SD_WPV_ was 0.5 mm [95% HDI: 0.42, 0.58] with a 100% posterior probability of exceeding the null.

For CON, the integrated estimate of ATE of VL MT was 0.4 mm [95% HDI: -0.25, 1; 95% HDPI: - 2.4, 3.24] with a 90.6% posterior probability of exceeding the null. On the participant level, ITE ranged from -0.84 mm to 1 mm with the 95% HDI of 16 out of 16 individuals suggesting estimates were compatible with the ATE. The resulting estimate of SD_IRV_ was 0.94 mm [95% HDI: 0.11, 1.7] with a 100% posterior probability of exceeding the null. The estimate of SD_WPV_ was 1 mm [95% HDI: 0.83, 1.14] with a 100% posterior probability of exceeding the null. These results can be visualized in the supplementary materials.

##### Reliability

For GEN, the average difference between phases in the ATE of VL CSA was -0.43 cm^2^ [95% HDI: - 2.06, 1.28; 95% HDPI: -9.85, 8.8] with a 68.82% posterior probability of exceeding the null. The highest density region of phase-specific training effects ranged from -2.62 cm^2^ to 15.15 cm^2^ in phase 1 and -3.03 cm^2^ to 14.76 cm^2^ in phase 2. These results can be visualized in Figure 7B. The resulting estimate of Pearson’s correlation coefficient was 0.7 [95% HDI: 0.46, 0.88] with a 100% posterior probability of exceeding the null. The estimate of the ICC was 0.83 [95% HDI: 0.65, 0.94] with a 100% posterior probability of exceeding the null.

For CON, the average difference between phases in the ATE of VL CSA was -1.47 cm^2^ [95% HDI: -4.1, 1.11; 95% HDPI: -19.99, 17.19]. The highest density region of phase-specific training effects ranged from -7.46 cm^2^ to 12.69 cm^2^ in phase 1 and -9 cm^2^ to 11.2 cm^2^ in phase 2. These results can be visualized in Figure 8B. The resulting estimate of Pearson’s correlation coefficient was 0.06 [95% HDI: -0.16, 0.45] with a 76.48% posterior probability of exceeding the null. The estimate of the ICC was 0 [95% HDI: 0, 0.54] with a 100% posterior probability of exceeding the null.

The results for VL MT largely mirrored the primary outcome. For GEN, the average difference between phases in the ATE of VL MT was 0.23 mm [95% HDI: -0.27, 0.71; 95% HDPI: -2.59, 3] with a 80.01% posterior probability of exceeding the null. The highest density region of phase-specific training effects ranged from -0.8 mm to 4.02 mm in phase 1 and -0.59 mm to 4.23 mm in phase 2. The resulting estimate of Pearson’s correlation coefficient was 0.64 [95% HDI: 0.36, 0.85] with a 99.99% posterior probability of exceeding the null. The estimate of the ICC was 0.81 [95% HDI: 0.56, 0.92] with a 100% posterior probability of exceeding the null.

For CON, the average difference between phases in the ATE of VL MT was -0.18 mm [95% HDI: - 1.02, 0.58; 95% HDPI: -5.79, 5.36]. The highest density region of phase-specific training effects ranged from -2.98 mm to 3.98 mm in phase 1 and -3.2 mm to 3.78 mm in phase 2. The resulting estimate of Pearson’s correlation coefficient was 0.28 [95% HDI: -0.05, 0.65] with a 94.98% posterior probability of exceeding the null. The estimate of the ICC was 0.47 [95% HDI: 0, 0.74] with a 100% posterior probability of exceeding the null. These results can be visualized in the supplementary materials.

##### Agreement

For GEN, the highest density region of differences between the ITE and the ATE for VL CSA ranged from -7.32 cm^2^ to 7.36 cm^2^. Alternatively, the highest density region of differences between the ITE with and without observation-level error ranged from -3.24 cm^2^ to 3.26 cm^2^. Consequently the limits of agreement extended from 3.23 cm^2^ [95% HDI: 2.75, 3.79] to -3.23 cm^2^ [95% HDI: -2.75, -3.79]. These results can be visualized in Figure 7C. The resulting estimate of lnVR was 0.77 [95% HDI: 0.23, 1.25] with a 99.75% posterior probability of exceeding the null. The estimate of DiV was 5.97 [95% HDI: 0.55, 25.03] with a 99.75% posterior probability of exceeding the null.

For CON, the highest density region of differences between the ITE and the ATE for VL CSA ranged from -3.98 cm^2^ to 3.99 cm^2^. Alternatively, the highest density region of differences between the ITE with and without observation-level error ranged from -6.43 cm^2^ to 6.53 cm^2^. Consequently the limits of agreement extended from 6.5 cm^2^ [95% HDI: 5.51, 7.58] to -6.5 cm^2^ [95% HDI: -5.51, -7.58]. These results can be visualized in Figure 8C. The resulting estimate of lnVR was -0.58 [95% HDI: -3.45, 0.41] with a 92.92% posterior probability of exceeding the null. The estimate of DiV was -9.58 [95% HDI: -15.06, 2.53] with a 92.92% posterior probability of exceeding the null.

The results for VL MT largely mirrored the primary outcome. For GEN, the highest density region of differences between the ITE and the ATE for VL MT ranged from -1.91 mm to 1.91 mm. Alternatively, the highest density region of differences between the ITE with and without observation-level error ranged from -0.97 mm to 0.97 mm. Consequently the limits of agreement extended from 0.99 mm [95% HDI: 0.82, 1.14] to -0.99 mm [95% HDI: -0.82, -1.14]. The resulting estimate of lnVR was 0.63 [95% HDI: 0.05, 1.12] with a 98.32% posterior probability of exceeding the null. The estimate of DiV was 0.32 [95% HDI: -0.05, 1.63] with a 98.32% posterior probability of exceeding the null.

For CON, the highest density region of differences between the ITE and the ATE for VL MT ranged from -2.08 mm to 2.09 mm. Alternatively, the highest density region of differences between the ITE with and without observation-level error ranged from -1.94 mm to 1.95 mm. Consequently the limits of agreement extended from 1.98 mm [95% HDI: 1.64, 2.27] to -1.98 mm [95% HDI: -1.64, -2.27]. The resulting estimate of lnVR was -0.06 [95% HDI: -1.32, 0.77] with a 58.23% posterior probability of exceeding the null. The estimate of DiV was -0.42 [95% HDI: - 1.22, 1.74] with a 58.23% posterior probability of exceeding the null. These results can be visualized in the supplementary materials.

### Strength

#### Test-Retest Reliability

The ICC for leg press 1RM was 0.98 [95% CI: 0.97, 0.99] with a SEM of 7.75 kg [95% CI: 6.21, 10.3] and CV of 4.68% [95% CI: 3.75, 6.23]. For knee extension MVIC, the ICC was 0.95 [95% CI: 0.92, 0.97] with a SEM of 4.33 kg [95% CI: 3.69, 5.24] and CV of 6.01% [95% CI: 5.11, 7.28]. These results can be visualized in Figure 9.

**Figure 9:**
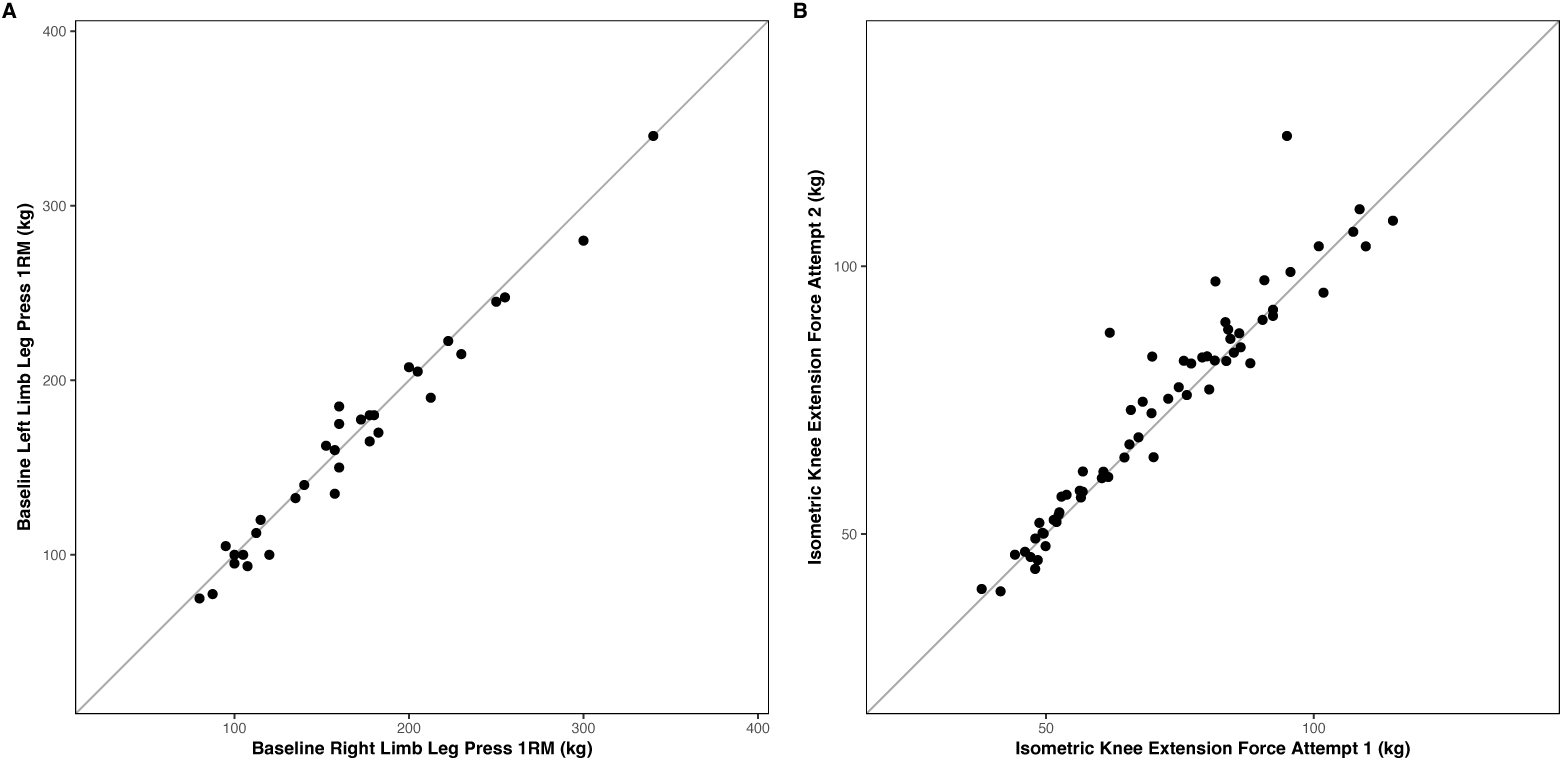
Test-retest reliability of A) leg press one-repetition maximum (1RM) and B) knee extension maximum voluntary isometric contraction (MVIC). The x-axis represents the value obtained at the first measurement occasion, while the y-axis represents the value obtained at the second occasion.

#### Gross Variability

Specifically for GEN, there was a range of -17.01 kg to 102.77 kg. When treating each phase independently, 30 participants demonstrated positive changes in leg press 1RM while 2 participants experienced negative change scores in leg press 1RM.

For CON, there was a range of -3.48 kg to 102.22 kg for HV conditions and -30.54 kg to 103.32 kg for LV conditions. When treating each phase independently, 19 participants demonstrated superior changes in the HV condition while 13 participants experienced superior change scores with the LV condition. These results are visualized in Figure 10.

**Figure 10:**
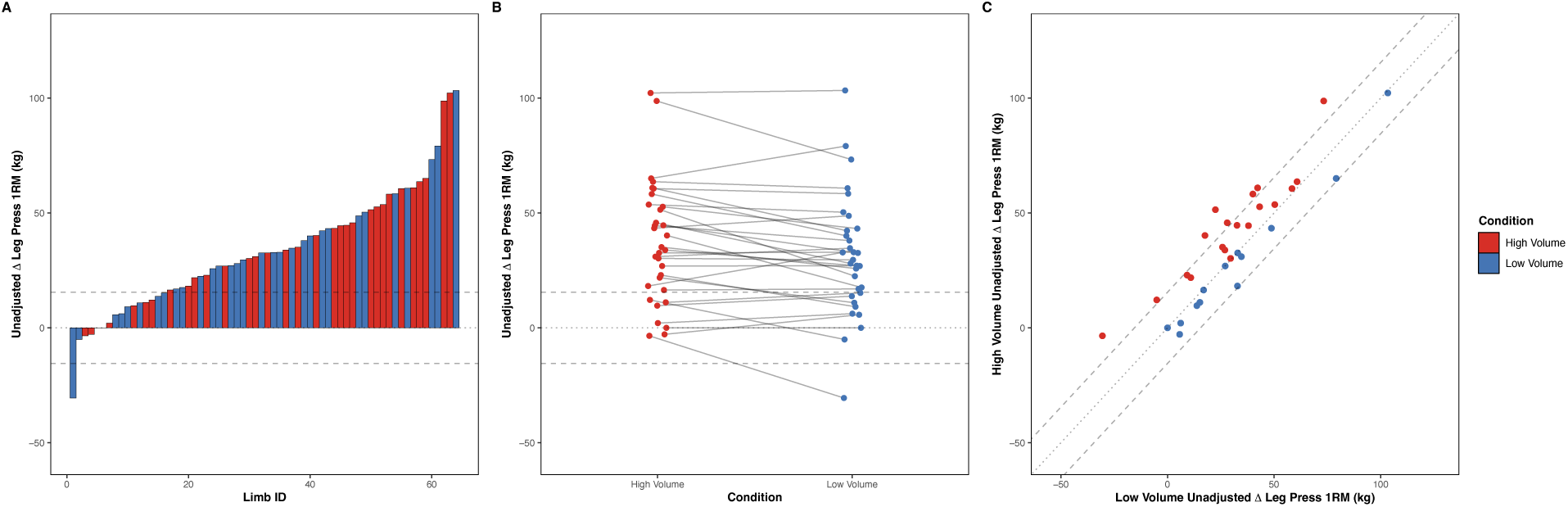
Gross variability of unadjusted change scores in leg press one-repetition maximum (1RM) following resistance training with high (red) or low (blue) weekly set volumes. The gray dashed lines represent the threshold derived from the estimate of measurement error (standard error of measurement (SEM) x 2) relative to a null change. A) waterfall plot in which each bar represents the unadjusted change score of a given limb, B) line plot in which the unadjusted change scores of each participant within a given phase is connected by a gray line, c) diagonal plot in which the x-axis represents the unadjusted change score of the low volume condition while the y-axis represents the high volume condition for a given participant. The dark diagonal line is the line of identity which represents a perfect correlation in the change scores between conditions. Red points are those who favor high volumes and blue points are those who favor low volumes.

The results for knee extension MVIC largely mirrored the primary outcome. Specifically for GEN, there was a range of -15.4 kg to 17.29 kg. When treating each phase independently, 16 participants demonstrated positive changes in knee extension MVIC while 16 participants experienced negative change scores in knee extension MVIC.

For CON, there was a range of -10.08 kg to 17.59 kg for HV conditions and -22.02 kg to 20.44 kg for LV conditions. When treating each phase independently, 20 participants demonstrated superior changes in the HV condition while 12 participants experienced superior change scores with the LV condition. These results can be visualized in the supplementary materials.

#### Naïve Estimates of Inter-individual Response Variation

##### Magnitude

For GEN, the naïve estimate of the ATE for leg press 1RM was 33.95 kg [95% CI: 24.91, 43; 95% PI: -14.41, 82.32]. On the participant level, ITE ranged from 7.87 kg to 69.86 kg with the 95% CI of 15 out of 16 individuals suggesting estimates were compatible with the ATE. These results can be visualized in Figure 11A. The resulting estimate of SD_IRV_ was 16.97 kg [95% CI: 11.29, 22.65] while SD_WPV_ was 27.01 kg [95% CI: 17.48, 36.55].

**Figure 11:**
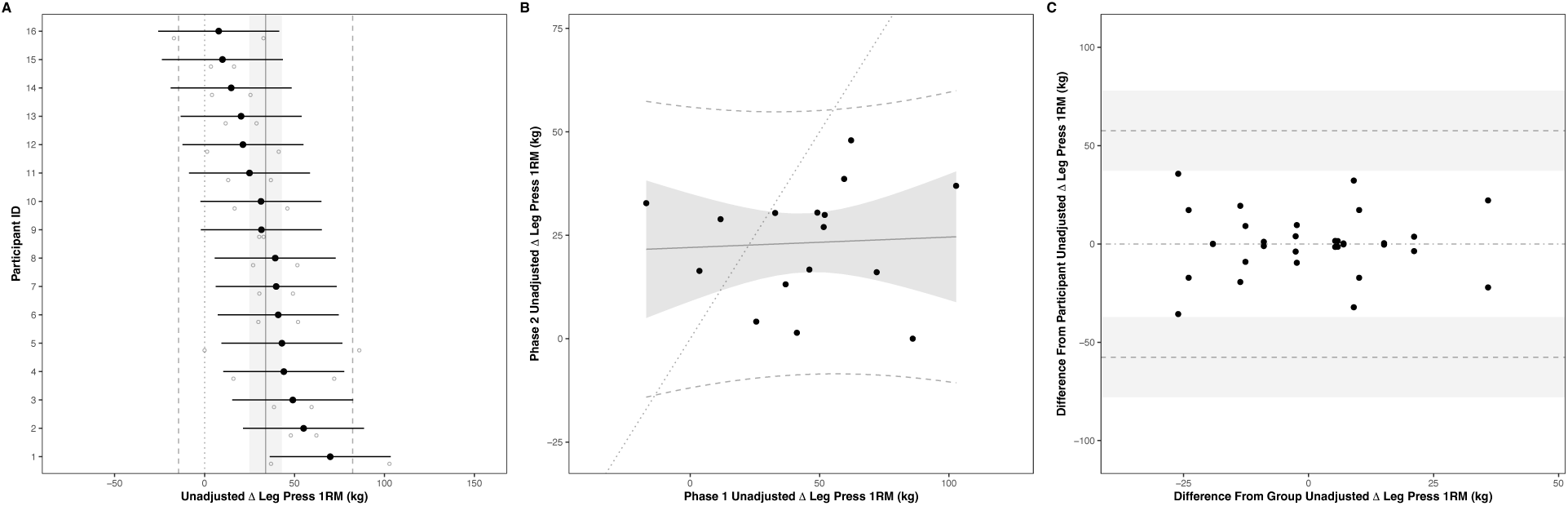
Naïve estimates of general (GEN) individual response variation (IRV) in leg press one-repetition maximum (1RM) following resistance training. The gray line shaded region and dashed lines represent the average training effect (ATE), 95% confidence, and prediction intervals, respectively. A) Forest plot of individual training effects (ITE). Black dots represent a given participants’ mean and uncertainty interval, while open circles represent the observed change score from each phase. B) Reliability of training effects between phases. Black dots represent the intersection of observed change scores from each phase. C) Agreement of training effects between phases. X-axis represents the deviation of ITE from the ATE, while the y-axis represents the deviation of the observed change score from each phase from their respective ITE.

For CON, the naïvely estimated ATE of leg press 1RM was 6.04 kg [95% CI: 2.27, 9.8; 95% PI: - 21.81, 33.89]. On the participant level, ITE ranged from -9.37 kg to 14.07 kg with the 95% CI of 16 out of 16 individuals suggesting estimates were compatible with the ATE. These results can be visualized in Figure 12A. The resulting estimate of SD_IRV_ was 7.07 kg [95% CI: 4.76, 9.38] while SD_WPV_ was 12.95 kg [95% CI: 9.67, 16.22].

**Figure 12:**
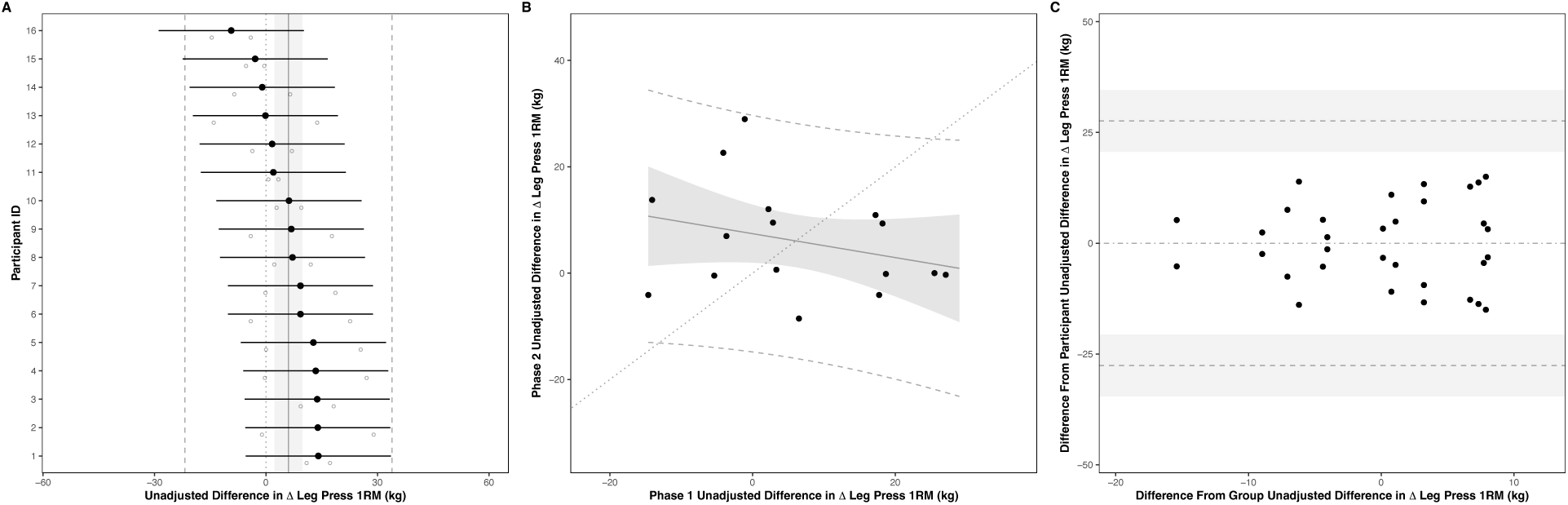
Naïve estimates of condition-specific (CON) individual response variation (IRV) in leg press one-repetition maximum (1RM) following resistance training. The gray line shaded region and dashed lines represent the average training effect (ATE), 95% confidence, and prediction intervals, respectively. A) Forest plot of individual training effects (ITE). Black dots represent a given participants’ mean and uncertainty interval, while open circles represent the observed change score from each phase. B) Reliability of training effects between phases. Black dots represent the intersection of observed change scores from each phase. C) Agreement of training effects between phases. X-axis represents the deviation of ITE from the ATE, while the y-axis represents the deviation of the observed change score from each phase from their respective ITE.

The results for knee extension MVIC largely mirrored the primary outcome. For GEN, the naïvely estimated ATE was 0.06 kg [95% CI: -2.02, 2.14; 95% PI: -14.85, 14.98]. On the participant level, ITE ranged from -5.27 kg to 6.69 kg with the 95% CI of 16 out of 16 individuals suggesting estimates were compatible with the ATE. The resulting estimate of SD_IRV_ was 3.91 kg [95% CI: 2.81, 5] while SD_WPV_ was 7.71 kg [95% CI: 4.76, 10.67].

For CON, the naïvely estimated ATE of knee extension MVIC was 1.57 kg [95% CI: -1.36, 4.5; 95% PI: -9.73, 12.86] indicating an advantage to the HV condition on the group-level. On the participant level, ITE ranged from -9.32 kg to 16.05 kg with the 95% CI of 14 out of 16 individuals suggesting estimates were compatible with the ATE. The resulting estimate of SD_IRV_ was 5.5 kg [95% CI: 2.67, 8.33] while SD_WPV_ was 5.19 kg [95% CI: 3.43, 6.96]. These results can be visualized in the supplementary materials.

##### Reliability

For GEN, the average difference between phases in the ATE of leg press 1RM was -21.58 kg [95% CI: -38.93, -4.23; 95% PI: -72.16, 29.01]. On the participant level, phase-specific training effects ranged from 18.66 kg to 80.65 kg in phase 1 and -2.92 kg to 59.07 kg in phase 2. These results can be visualized in Figure 11B. The resulting estimate of Pearson’s correlation coefficient was 0.05 [95% CI: -0.53, 0.64] while the ICC was 0.28 [95% CI: 0.11, 0.45].

For CON, the average difference between phases in the ATE of leg press 1RM was 0.05 kg [95% CI: -10.03, 10.12; 95% PI: -29.33, 29.42]. On the participant level, phase-specific training effects ranged from -14.62 kg to 27.06 kg in phase 1 and -8.54 kg to 28.96 kg in phase 2. These results can be visualized in Figure 12B. The resulting estimate of Pearson’s correlation coefficient was - 0.29 [95% CI: -0.74, 0.15] while the ICC was 0.23 [95% CI: 0.07, 0.39].

The results for knee extension MVIC largely mirrored the primary outcome. For GEN, the average difference between phases in the ATE of knee extension MVIC was 4.79 kg [95% CI: - 0.6, 10.18; 95% PI: -10.93, 20.51]. On the participant level, phase-specific training effects ranged from -7.66 kg to 4.3 kg in phase 1 and -2.87 kg to 9.09 kg in phase 2. The resulting estimate of Pearson’s correlation coefficient was -0.26 [95% CI: -0.74, 0.23] while the ICC was 0.2 [95% CI: 0.06, 0.35].

For CON, the average difference between phases in the ATE of knee extension MVIC was -1.25 kg [95% CI: -5.23, 2.74; 95% PI: -12.86, 10.37]. On the participant level, phase-specific training effects ranged from -12.34 kg to 16.29 kg in phase 1 and -6.3 kg to 15.8 kg in phase 2. The resulting estimate of Pearson’s correlation coefficient was 0.42 [95% CI: -0.34, 1.18] while the ICC was 0.53 [95% CI: 0.2, 0.86]. These results can be visualized in the supplementary materials.

##### Agreement

For GEN, the differences between the ITE and the ATE for leg press 1RM ranged from -35.67 kg to 35.67 kg. Alternatively, the difference between the ITE with and without observation-level error ranged from -35.67 kg to 35.67 kg. Consequently the limits of agreement extended from 57.58 kg [95% CI: 37.26, 77.9] to -57.58 kg [95% CI: -37.26, -77.9]. These results can be visualized in Figure 11C. The resulting estimate of lnVR was -0.93 [95% CI: -1.82, -0.04], while DiV was -441.86 [95% CI: -935.41, 51.7].

For CON, the differences between the ITE and the ATE for leg press 1RM ranged from -15.01 kg to 15.01 kg. Alternatively, the difference between the ITE with and without observation-level error ranged from -15.01 kg to 15.01 kg. Consequently the limits of agreement extended from 27.59 kg [95% CI: 20.62, 34.57] to – 27.59 kg [95% CI: -20.62, -34.57]. These results can be visualized in Figure 12C. The resulting estimate of lnVR was -1.21 [95% CI: -2.13, -0.29], while DiV was -117.65 [95% CI: -208.89, -26.41].

The results for knee extension MVIC largely mirrored the primary outcome. For GEN, the differences between the ITE and the ATE for knee extension MVIC ranged from -9.9 kg to 9.9 kg. Alternatively, the difference between the ITE with and without observation-level error ranged from -9.9 kg to 9.9 kg. Consequently the limits of agreement extended from 16.44 kg [95% CI: 10.14, 22.74] to -16.44 kg [95% CI: -10.14, -22.74]. The resulting estimate of lnVR was -1.36 [95% CI: -2.25, -0.47], while DiV was -44.24 [95% CI: -88.46, -0.01].

For CON, the differences between the ITE and the ATE for knee extension MVIC ranged from - 7.12 kg to 7.12 kg. Alternatively, the difference between the ITE with and without observation-level error ranged from -7.12 kg to 7.12 kg. Consequently the limits of agreement extended from 11.07 kg [95% CI: 7.31, 14.83] to -11.07 kg [95% CI: -7.31, -14.83]. The resulting estimate of lnVR was 0.11 [95% CI: -1.34, 1.57], while DiV was 3.24 [95% CI: -33.04, 39.52]. These results can be visualized in the supplementary materials.

#### Integrated Estimates of Inter-individual Response Variation

##### Magnitude

For GEN, the integrated estimate of ATE of leg press 1RM was 35.04 kg [95% HDI: 24.67, 45.36; 95% HDPI: 9.84, 60.55] with a 100% posterior probability of exceeding the null. On the participant level, ITE ranged from 19.25 kg to 59.58 kg with the 95% HDI of 15 out of 16 individuals suggesting estimates were compatible with the ATE. These results can be visualized in Figure 13A and 14A, respectively. The resulting estimate of SD_IRV_ was 14.1 kg [95% HDI: 5.61, 27.59] with a 100% posterior probability of exceeding the null. The estimate of SD_WPV_ was 8.2 kg [95% HDI: 6.82, 9.84] with a 100% posterior probability of exceeding the null.

**Figure 13:**
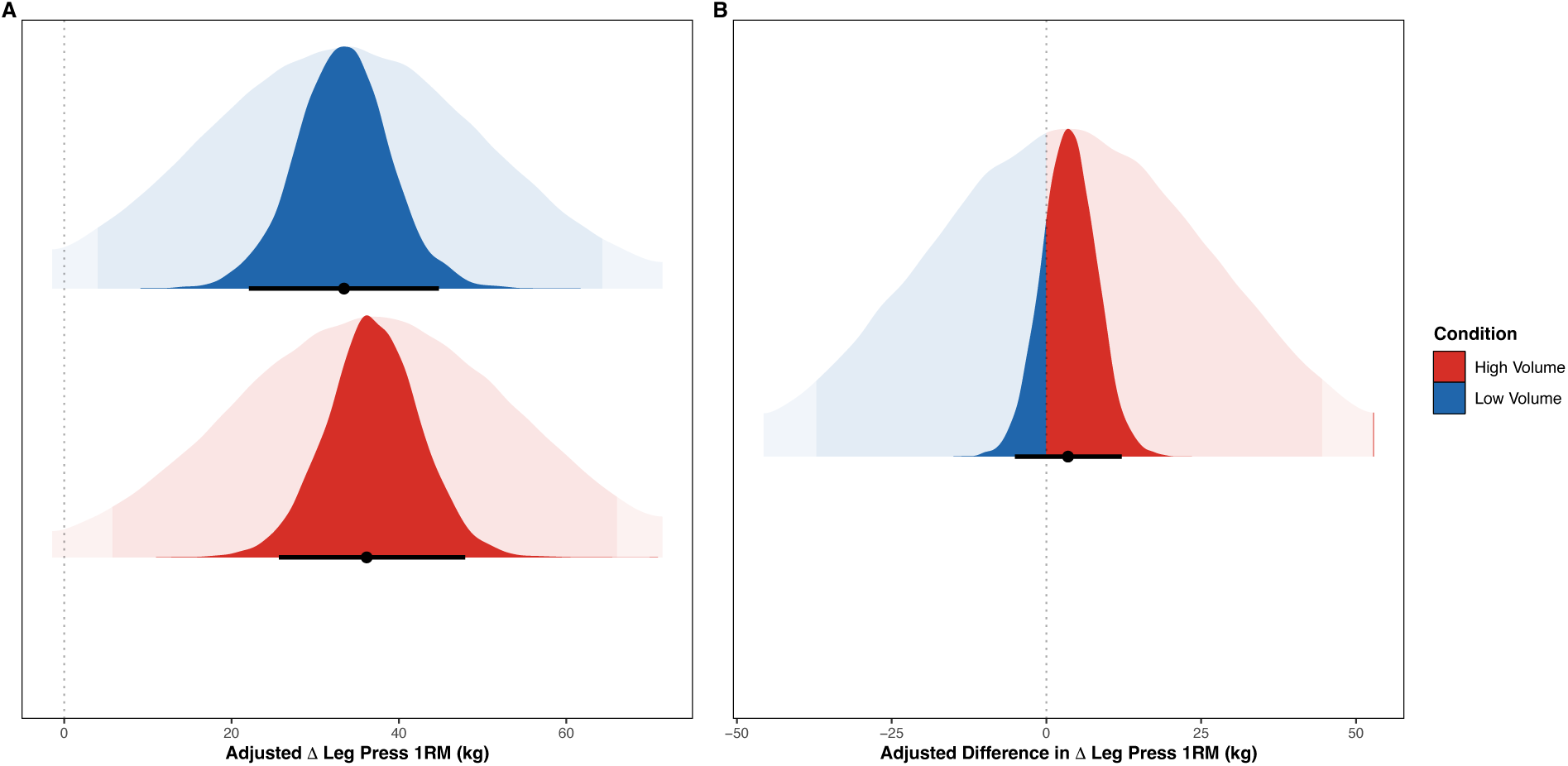
Adjusted average training effects (ATE) for changes in leg press one-repetition maximum (1RM) A) within, and B) between conditions. Black dots and intervals represent the mode and the 95% highest density intervals (HDI) of the posterior distribution while the faded densities in the background represent the draws from the posterior predictive distribution, where the darker region represents the 95% prediction interval. For within-condition effects, the red distributions represents high volumes (HV) and the blue distributions represents low volumes (LV). For the between-condition effect, the red portion of the distributions represents the posterior draws that favor HV while the blue portion of the distributions favor LV.

**Figure 14:**
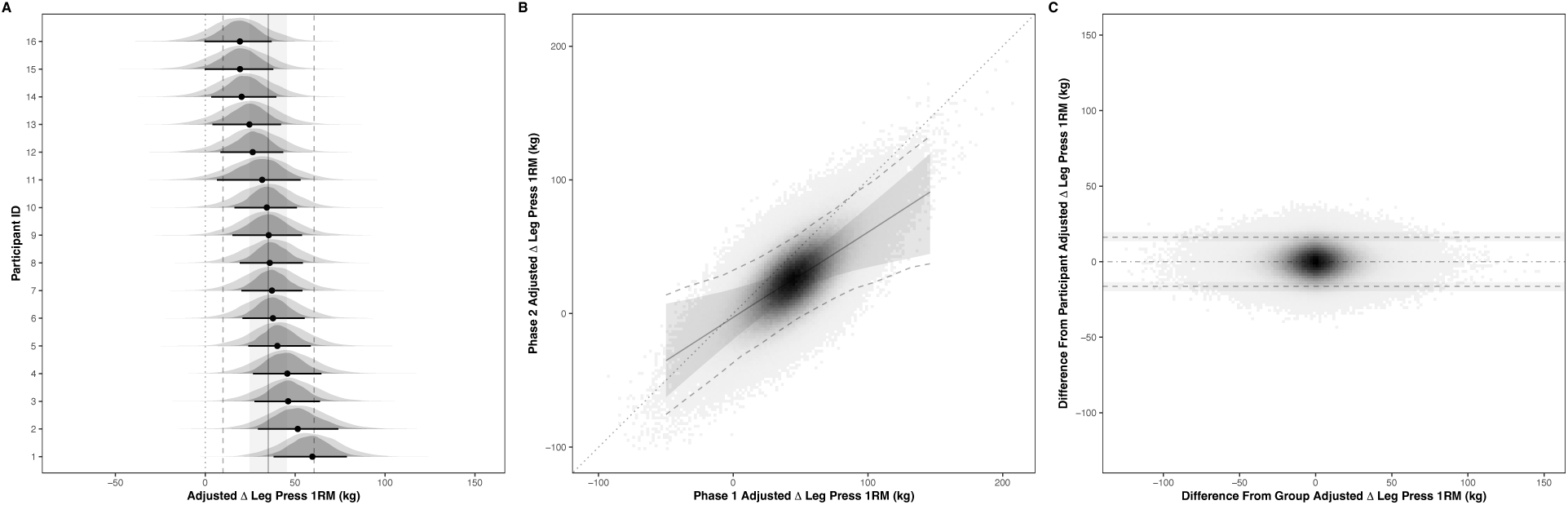
Integrated estimates of general (GEN) of individual response variation (IRV) in leg press one-repetition maximum (1RM) following resistance training. The gray line shaded region and dashed lines represent the average training effect (ATE), 95% highest density, and prediction intervals, respectively. A) Forest plot of the posterior distributions of individual training effects (ITE). Black dots and uncertainty intervals represent the participants’ mode and uncertainty interval while the faded densities represent the posterior predictive distribution. B) Reliability of training effects between phases. Colored bins represent the intersection posterior predictions of training effects of each phase from simulated participants. C) Agreement of training effects between phases. X-axis represents the deviation of simulated ITE from the ATE, while the y-axis represents the deviation of the simulated posterior predictions from each phase from their respective ITE.

For CON, the integrated estimate of ATE of leg press 1RM was 3.48 kg [95% HDI: -5.1, 12.15; 95% HDPI: -43.28, 50.69] with a 80.01% posterior probability of exceeding the null. On the participant level, ITE ranged from 1.71 kg to 4.47 kg with the 95% HDI of 16 out of 16 individuals suggesting estimates were compatible with the ATE. These results can be visualized in Figures 13B and 15A, respectively. The resulting estimate of SD_IRV_ was 1.45 kg [95% HDI: 0, 11.54] with a 100% posterior probability of exceeding the null. The estimate of SD_WPV_ was 16.31 kg [95% HDI: 13.56, 19.59] with a 100% posterior probability of exceeding the null.

**Figure 15:**
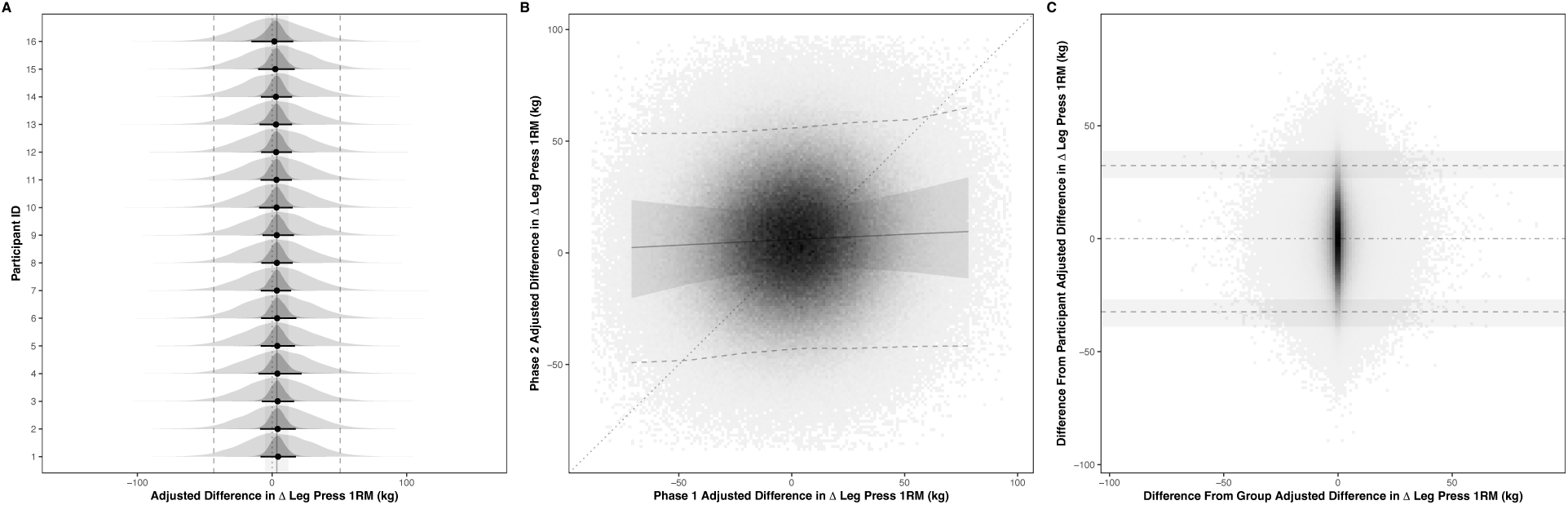
Integrated estimates of condition-specific (CON) individual response variation (IRV) in leg press one-repetition maximum (1RM) following resistance training. The gray line shaded region and dashed lines represent the average training effect (ATE), 95% highest density, and prediction intervals, respectively. A) Forest plot of the posterior distributions of individual training effects (ITE). Black dots and uncertainty intervals represent the participants’ mode and uncertainty interval while the faded densities represent the posterior predictive distribution. B) Reliability of training effects between phases. Colored bins represent the intersection posterior predictions of training effects of each phase from simulated participants. C) Agreement of training effects between phases. X-axis represents the deviation of simulated ITE from the ATE, while the y-axis represents the deviation of the simulated posterior predictions from each phase from their respective ITE.

The results for knee extension MVIC largely mirrored the primary outcome. For GEN, the integrated estimate of ATE of knee extension MVIC was 0.49 kg [95% HDI: -2.59, 3.35; 95% HDPI: -10.47, 11.34] with a 62.36% posterior probability of exceeding the null. On the participant level, ITE ranged from -0.13 kg to 1.3 kg with the 95% HDI of 16 out of 16 individuals suggesting estimates were compatible with the ATE. The resulting estimate of SD_IRV_ was 1.48 kg [95% HDI: 0, 5.65] with a 100% posterior probability of exceeding the null. The estimate of SD_WPV_ was 3.71 kg [95% HDI: 3.09, 4.45] with a 100% posterior probability of exceeding the null.

For CON, the integrated estimate of ATE of knee extension MVIC was 0.79 kg [95% HDI: -3.83, 5.69; 95% HDPI: -20.48, 22.34] with a 64.72% posterior probability of exceeding the null. On the participant level, ITE ranged from -1.05 kg to 2.28 kg with the 95% HDI of 16 out of 16 individuals suggesting estimates were compatible with the ATE. The resulting estimate of SD_IRV_ was 3.41 kg [95% HDI: 0, 10.34] with a 100% posterior probability of exceeding the null. The estimate of SD_WPV_ was 7.47 kg [95% HDI: 6.14, 8.87] with a 100% posterior probability of exceeding the null. These results can be visualized in the supplementary materials.

##### Reliability

For GEN, the average difference between phases in the ATE of leg press 1RM was -18.87 kg [95% HDI: -30.72, -7.25; 95% HDPI: -67.99, 27.79] with a 99.88% posterior probability of exceeding the null. The highest density region of phase-specific training effects ranged from 1.25 kg to 87.81 kg in phase 1 and -17.89 kg to 68.89 kg in phase 2. These results can be visualized in Figure 14B. The resulting estimate of Pearson’s correlation coefficient was 0.67 [95% HDI: 0.27, 0.9] with a 99.72% posterior probability of exceeding the null. The estimate of the ICC was 0.85 [95% HDI: 0.47, 0.96] with a 100% posterior probability of exceeding the null.

For CON, the average difference between phases in the ATE of leg press 1RM was 5.12 kg [95% HDI: -11.15, 21.47; 95% HDPI: -89.35, 98.91]. The highest density region of phase-specific training effects ranged from -47.85 kg to 50.28 kg in phase 1 and -43.31 kg to 55.16 kg in phase 2. These results can be visualized in Figure 15B. The resulting estimate of Pearson’s correlation coefficient was 0.04 [95% HDI: -0.19, 0.29] with a 65.72% posterior probability of exceeding the null. The estimate of the ICC was 0 [95% HDI: 0, 0.33] with a 100% posterior probability of exceeding the null.

The results for knee extension MVIC largely mirrored the primary outcome. For GEN, the average difference between phases in the ATE of knee extension MVIC was 6.48 kg [95% HDI: 1.08, 11.54; 95% HDPI: -15.25, 27.94] with a 99.04% posterior probability of exceeding the null. The highest density region of phase-specific training effects ranged from -15.37 kg to 9.88 kg in phase 1 and -9.11 kg to 16.35 kg in phase 2. The resulting estimate of Pearson’s correlation coefficient was 0.11 [95% HDI: -0.16, 0.59] with a 83.15% posterior probability of exceeding the null. The estimate of the ICC was 0 [95% HDI: 0, 0.7] with a 100% posterior probability of exceeding the null.

For CON, the average difference between phases in the ATE of knee extension MVIC was 5.2 kg [95% HDI: -2.3, 13.08; 95% HDPI: -37.41, 48.02]. The highest density region of phase-specific training effects ranged from -26.29 kg to 22.76 kg in phase 1 and -21.18 kg to 28.06 kg in phase 2. The resulting estimate of Pearson’s correlation coefficient was 0.12 [95% HDI: -0.15, 0.55] with a 84.61% posterior probability of exceeding the null. The estimate of the ICC was 0 [95% HDI: 0, 0.67] with a 100% posterior probability of exceeding the null. These results can be visualized in the supplementary materials.

##### Agreement

For GEN, the highest density region of differences between the ITE and the ATE for leg press 1RM ranged from -35.07 kg to 35.25 kg. Alternatively, the highest density region of differences between the ITE with and without observation-level error ranged from -16.37 kg to 16.38 kg. Consequently the limits of agreement extended from 16.27 kg [95% HDI: 13.54, 19.53] to -16.27 kg [95% HDI: -13.54, -19.53]. These results can be visualized in Figure 14C. The resulting estimate of lnVR was 0.71 [95% HDI: -0.15, 1.36] with a 94.08% posterior probability of exceeding the null. The estimate of DiV was 105.21 [95% HDI: -80.29, 628.68] with a 94.08% posterior probability of exceeding the null.

For CON, the highest density region of differences between the ITE and the ATE for leg press 1RM ranged from -12.38 kg to 12.46 kg. Alternatively, the highest density region of differences between the ITE with and without observation-level error ranged from -32.9 kg to 32.63 kg. Consequently the limits of agreement extended from 32.36 kg [95% HDI: 26.9, 38.87] to -32.36 kg [95% HDI: -26.9, -38.87]. These results can be visualized in Figure 15C. The resulting estimate of lnVR was -1.33 [95% HDI: -4, -0.02] with a 99.3% posterior probability of exceeding the null. The estimate of DiV was -247.57 [95% HDI: -383.32, -100.45] with a 99.3% posterior probability of exceeding the null.

The results for knee extension MVIC largely mirrored the primary outcome. For GEN, the highest density region of differences between the ITE and the ATE for knee extension MVIC ranged from -6.46 kg to 6.43 kg. Alternatively, the highest density region of differences between the ITE with and without observation-level error ranged from -7.37 kg to 7.44 kg. Consequently the limits of agreement extended from 7.37 kg [95% HDI: 6.13, 8.82] to -7.37 kg [95% HDI: -6.13, -8.82]. The resulting estimate of lnVR was -0.17 [95% HDI: -3.02, 0.84] with a 78.79% posterior probability of exceeding the null. The estimate of DiV was -12.16 [95% HDI: - 20.43, 19.93] with a 78.79% posterior probability of exceeding the null.

For CON, the highest density region of differences between the ITE and the ATE for knee extension MVIC ranged from -12.19 kg to 12.25 kg. Alternatively, the highest density region of differences between the ITE with and without observation-level error ranged from -14.81 kg to 14.82 kg. Consequently the limits of agreement extended from 14.82 kg [95% HDI: 12.19, 17.6] to -14.82 kg [95% HDI: -12.19, -17.6]. The resulting estimate of lnVR was -0.15 [95% HDI: -2.81, 0.68] with a 80.8% posterior probability of exceeding the null. The estimate of DiV was -49.75 [95% HDI: -80.54, 58.04] with a 80.8% posterior probability of exceeding the null. These results can be visualized in the supplementary materials.

## DISCUSSION

This study employed a novel replicated within-participant unilateral design to investigate both GEN and CON IRV following RT in recreationally trained participants. Our hypotheses for this experiment were largely confirmed. Despite observing clear gross variability across all outcomes, we failed to reveal irrefutable evidence of true IRV across our multi-stage statistical approach. However, the evidence was considerably stronger for GEN IRV when integrating information across participants, likely reflecting the greater statistical power and precision of these methods. This pattern suggests that there may be some detectable variation in how individuals respond to RT independent of the specific manipulation of weekly set volume. On the group-level, higher weekly set volumes (16 vs. 8 sets per week) led to a detectable advantage for muscle hypertrophy but not maximal strength outcomes.

### Group-level Findings

Our group-level findings align well with recent meta-analytic evidence from Pelland et al. (Pelland *et al*., 2024). For muscle hypertrophy, higher weekly set volumes demonstrated a detectable advantage, with an ATE of VL CSA of 1.8 cm^2^ [95% HDI: 0.29, 3.41] and a 98.77% posterior probability of an advantage to HV. In contrast, maximal strength outcomes showed no detectable differences between conditions, with an ATE of leg press 1RM of 3.48 kg [95% HDI: - 5.1, 12.15] and a 80.01% posterior probability of an advantage to HV.

These results fit well with the non-linear dose-response relationships reported in the aforementioned meta-analysis. While the dose-response relationship between weekly set volume and muscle hypertrophy shows clear diminishing returns, the difference between predictions at 8 and 16 sets was still notable (2.01% [95% CrI: 1.20, 2.77]). Alternatively, the dose-response relationship between weekly set volume and maximal strength outcomes exhibited more aggressive attenuation from 8 to 16 sets with differences likely too small to be detected in the present study (0.77% [95% CrI: 0.58, 0.94]). These results support the notion that weekly training volume exerts greater influence on muscle hypertrophy than maximal strength outcomes, but this interpretation requires important context since the present training protocol was not specifically designed to optimize maximal strength gain.

Our study design prioritized the ability to compare our results to previous studies that are commonly used as evidence of IRV to RT with different weekly set volumes, rather than optimizing the training protocol for maximal strength development (Damas *et al*., 2019; Hammarström *et al*., 2020; Lixandrão *et al*., 2024). We utilized loads corresponding to 10-14 RM, that are less specific to the strength assessment (i.e., 1RM). Given that the specificity of load strongly predicts maximal strength gains (Swinton *et al*., 2024), this design choice may have substantially influenced our results. Notably, Pelland et al. reported similar training characteristics across the included studies, with an average of 9.85 ± 3.19 repetitions per set, suggesting comparable limitations in task specificity across the broader literature. Had our intervention employed training with greater specificity to the maximal strength assessment, there might have been differential effects of weekly set volume. These findings reinforce the notion that strength and hypertrophy adaptations are optimized through distinct organizations of training variables, at least within the typical duration of training studies (i.e., ∼8-12 weeks) (Refalo *et al*., 2021; Swinton *et al*., 2024; Robinson *et al*., 2024b; Pelland *et al*., 2024; Remmert *et al*., 2025).

### Individual-level Findings

Our individual-level findings demonstrated substantial gross variability across all outcomes, mirroring previous research often cited as evidence for IRV following RT with different weekly set volumes. However, despite this apparent heterogeneity, we failed to reveal irrefutable evidence of IRV across our multi-stage statistical analyses. When appropriately decomposing the confounding sources of variation, the observed gross variability was largely attributable to WPV rather than true inter-individual differences (i.e., IRV). This finding represents the primary contribution of our study to the overall body of research. Specifically, it demonstrates how IRV must be investigated with appropriate study design and statistical approaches to obtain valid inferences and unbiased estimates of the target causal effects.

While our study has multiple limitations, our results provide stronger evidence for GEN IRV compared to CON, particularly with integrated statistical methods. This pattern suggests that individuals may exhibit some true response variation following RT independent of the specific manipulation of weekly set volume. Specifically, the integrated correlation coefficients (i.e., Pearson’s r) ranged from 0.68 to 0.71 for GEN, while CON suggested much weaker relationships with a range of 0.04 to 0.06. Similarly, the integrated multiplicative contrasts (i.e., lnVR) between SD_IRV_ and SD_WPV_ ranged from 0.71 to 0.77 for GEN, while CON failed to reveal that the IRV was meaningful with a range of -1.33 to -0.58. These findings may reflect that, due to unclear factor(s), some individuals respond more favorably to RT generally, but these factors did not result in preferential responses to a given weekly set volume.

Our findings with respect to different weekly set volumes align with growing evidence across the domains of medicine, nutrition science, and exercise physiology, demonstrating that observed gross variability rarely represents real or meaningful inter-individual differences (Williamson *et al*., 2018; Kelley *et al*., 2021, 2023, 2024; Esteves *et al*., 2021; Kelley *et al*., 2022; Bonafiglia *et al*., 2022; Steele *et al*., 2023). The preponderance of evidence indicates that the apparent heterogeneity in intervention effects are typically best explained by measurement error, sampling variance, biological variability, and other confounding sources of within-participant variation rather than true differential responses to interventions between individuals. To be clear, these results do not suggest that the magnitude of true IRV is known to be zero. Rather, they indicate that the magnitude of true IRV, combined with the practical challenges of accurately estimating its magnitude with the available sample sizes, rarely justifies the additional complexity to improve our predictions and subsequent recommendations.

While this study is a small contribution to the broader literature rigorously investigating IRV in RT, it provides a clear example where intuition and appropriate inference do not align. Our results largely support the broader literature suggesting that population-based recommendations remain appropriate for most RT applications. Thus, the findings of the present study are only unique through perception, and in reality simply provide additional support to an under-appreciated sentiment across other domains of research.

### Limitations

Several limitations warrant consideration when interpreting our findings. First, we experienced some participant attrition, with two participants unable to complete post-testing in the second phase (one dropout, one training-related injury) and one participant unable to complete all angles of isometric strength testing, resulting in missing data that may have influenced our estimates.

Second, it is possible that the recruitment of participants for RT studies results in an artificially homogeneous sample relative to the broader population engaging in RT (i.e., selection bias). However, longitudinal analyses of the open powerlifting dataset (Latella *et al*., 2023), which is absent these constraints, demonstrate remarkably similar variance component ratios, with the variation explained by the random slopes of time (i.e., IRV) remaining modest compared to that explained by the random intercepts and remaining WPV. This convergence suggests that selection bias may not prevent our results from generalizing beyond our sample.

Third, the duration of the training phases in the present study (i.e., ∼11 weeks) might be considered insufficient for divergent responses, and therefore variation, to manifest. Yet, longitudinal analyses following individuals performing low-volume RT for up to ∼6 years also revealed similar variance component ratios (Steele *et al*., 2022), with the variation explained by the random slopes of time (i.e., IRV) remaining modest compared to that explained by the random intercepts and remaining WPV. These long term observations lend support to the idea that our estimates of IRV were not biased due to intervention length.

Fourth, the present study featured a limited sample size (i.e., n = 16, with 2 phases per participant) that likely compromised statistical power. However, recent meta-analyses (Williamson *et al*., 2018; Kelley *et al*., 2021, 2023, 2024; Esteves *et al*., 2021; Kelley *et al*., 2022; Bonafiglia *et al*., 2022; Steele *et al*., 2023), which pool the results of many studies to gain statistical power, demonstrate similar findings when comparing the variances between non-training controls and exercise conditions. The consistency with studies in which statistical power is comparatively less of a concern lends further support to the stability of our conclusions.

Fifth, our crossover design relies on assumptions that may not fully hold. Despite our 6-8 week washout period, we cannot definitively rule out carry-over effects, though our statistical models attempted to account for phase-specific differences. The absence of substantial phase-specific differences provides some reassurance, but this limitation remains inherent to crossover designs and the estimates thereof. Moreover, due to the novelty of our study design, it is not fully understood how the limb-centric design elements (e.g., cross-education effect, sequence allocation, etc.) influence the ability to estimate the target effects compared to a standard replicated crossover design.

Moreover, due to the lack of a time-matched non-training control group, we must make the assumption that changes in our primary outcomes are indeed caused by the training intervention rather than other confounding sources of error (e.g., measurement drift). While this is a limitation, the ability to obtain unbiased estimates of causal effects in the absence of a time-matched non-training control group seems to adequate in RT research (Steele *et al*., 2023). Specifically, the relationship between controlled (i.e., *Δ* RT - *Δ* Control) and uncontrolled (i.e., *Δ* RT) is nearly one to one (*β* = 0.90 to 0.95) (https://osf.io/nkwu8).

Finally, our investigation focused specifically on weekly set volume, and these results may not generalize to all training variables. Other factors such as exercise selection, repetition ranges, or training frequency might demonstrate different patterns of IRV. However, given the breadth of topics in which IRV has been shown to be modest compared to the remaining WPV (e.g., intervention-induced changes in body mass, body composition, cardiorespiratory fitness, waist circumference, blood pressure, muscle size, maximal strength, and time to exhaustion following beta-alanine supplementation), it may be reasonable to assume generalizability as the null hypothesis.

Critically, the limitations related to selection bias, intervention duration, and statistical power apply equally to studies often cited as evidence for meaningful IRV in RT. All previous investigations employed designs without replication, preventing separation of IRV from WPV. Our replicated within-participant design, despite its limitations, represents a methodological advance for investigating these questions.

### Practical Applications

This study demonstrates that seemingly intuitive interpretations of variability in training outcomes often lead to inappropriate inferences. The observation of gross variability does not necessarily indicate true inter-individual differences. Researchers must recognize that studies designed to investigate IRV require fundamentally different approaches than those examining average training effects.

Scientific studies typically pursue one of three inferential goals: to describe phenomena, predict outcomes, or explain causal effects. Most randomized trials aim to do the latter, but researchers must clearly specify whether they seek to investigate the expected magnitude of effects (i.e., mean) or the variance thereof. This distinction profoundly influences study design, statistical analysis, and required resources. For instance, detecting causal effects related to IRV would likely require much larger sample sizes than what is necessary for group-level outcomes, far exceeding what is typical in RT studies (Senn, 2017).

If investigating causal effects of IRV is the goal, it is paramount for researchers to understand the complete chain of evidence required to support personalized training recommendations (Figure 16). Most of the evidence in this area, including the present study, address the early links in this chain; revealing that despite clear gross variability, the evidence for reliable and meaningful IRV remains limited.

**Figure 16:**
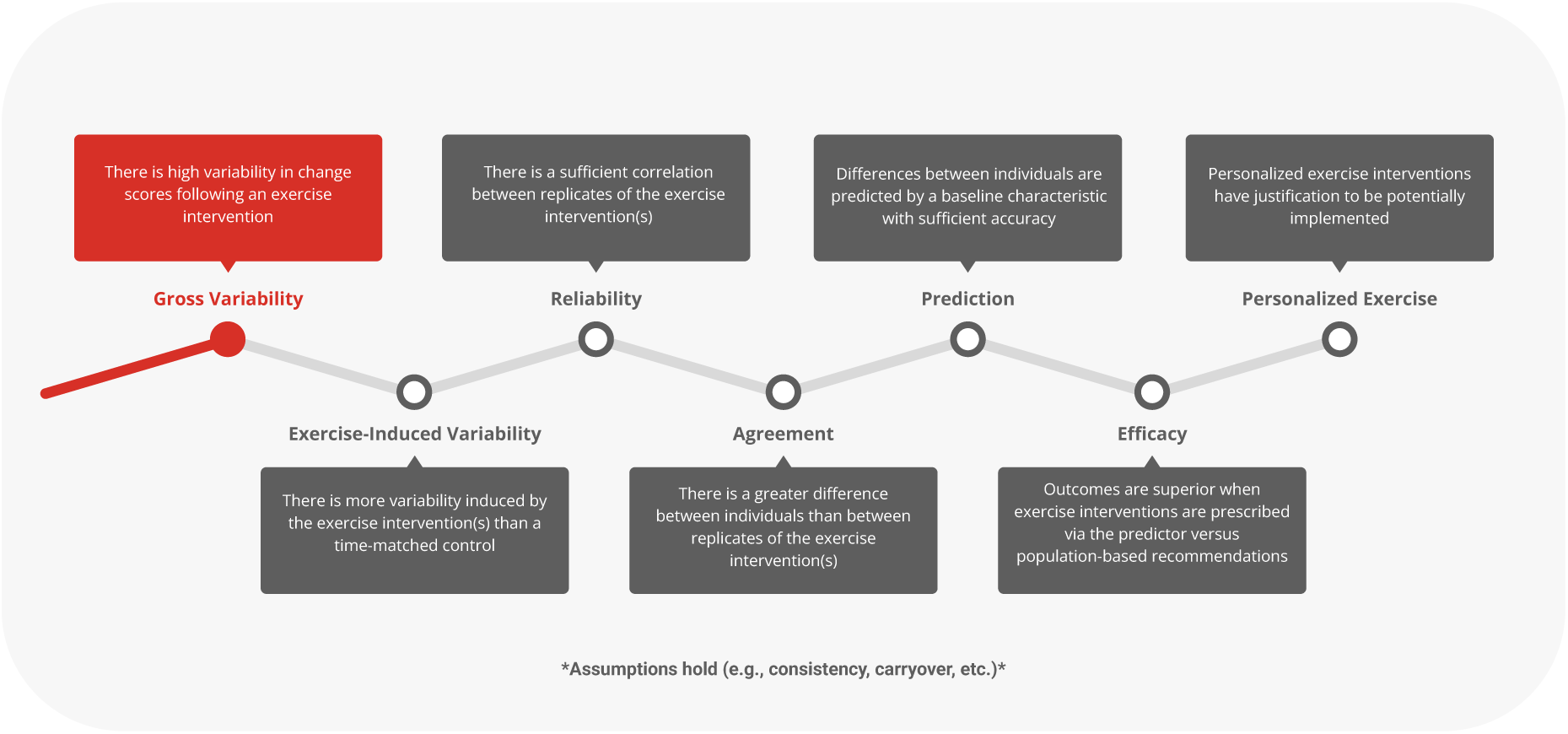
Conceptual map of the chain of evidence required to support personalized training recommendations.

It is crucial to clarify that this study does not argue against the individualization of RT programs. Rather, it suggests that the grounds on which individualization is based on may require further scrutiny. Specifically, individualization causally linked to specific training variables (e.g., volume) has limited empirical support. Our findings indicate that apparent differences in how individuals respond to 8 versus 16 weekly sets likely reflect the confounding sources of within-participant variation rather than stable, predictable characteristics. Instead, practitioners should consider individualizing programs predominantly based on factors external to the organization of the training intervention itself (e.g., tolerance, preferences, adherence, logistical constraints, prioritization, etc.).

For practitioners who remain unconvinced that IRV is negligible relative to the remaining WPV, the foundational concepts from this study design still apply. Any observed variability in training outcomes must be carefully delineated from the other sources of variation before concluding that a specific approach benefits a particular individual.

At the individual level, this presents substantial challenges due to the logarithmic nature of RT adaptations and the complex influence of the higher order effects of previous training history (Steele *et al*., 2022; Latella *et al*., 2023). Without borrowing information from a larger sample or repeated exposures to different training conditions, it is potentially impractical in real-world settings to distinguish signal from noise with a high degree of precision. Based on the current evidence, recommendations derived from population-level principles that are then further modified to meet the constraints of the individual are likely the best sources of information for program design.

### Conclusion

This study demonstrates the study design, statistical analyses, and resources necessary to appropriately investigate IRV in RT. By employing a replicated within-participant unilateral design, we separated confounding sources of variation that typically obscure the interpretation of training effects. Our findings clearly illustrate that gross variability in training outcomes does not necessarily indicate true inter-individual differences; a distinction critical for both research and practice. Despite meaningful limitations, our results suggest stronger evidence of true variation between individuals in their response to RT independent of the specific manipulation of weekly set volume (i.e., GEN IRV). Alternatively, we failed to reveal evidence that an individual’s response to a given weekly set volume is a stable, predictable characteristic (i.e., CON IRV). At the group level, our findings suggest higher weekly training volumes of moderate-load RT conferred detectable benefits for muscle hypertrophy but not maximal strength. These differential outcomes underscore the importance of aligning training variables with the specific outcome of interest.

## DEFINITIONS

RT: Resistance training
IRV: Inter-individual response variation
WPV: Within-participant variation
GEN: General inter-individual response variation
CON: Condition-specific inter-individual response variation
ATE: Average training effect
ITE: Individual training effects
SD_IRV_: Standard deviation of inter-individual response variation
SD_WPV_: Standard deviation of within-participant variation
LV: Low volume condition (i.e., 6-8 sets per week)
HV: High volume condition (i.e., 12-16 sets per week)
VL: Vastus lateralis
CSA: Cross-sectional area
1RM: One-repetition maximum
MT: Muscle thickness
MVIC: Maximal voluntary isometric contraction
RIR: Repetitions in reserve
MPV: Mean propulsive velocity
sRPE: Session rating of perceived exertion
ICC: Intraclass correlation coefficient
SEM: Standard error of measurement
CV: Coefficient of variation
CI: Confidence interval
PI: Prediction interval
lnVR: Log variability ratio
DiV: Difference in variability
LOA: Limits of agreement
ANCOVA: Analysis of covariance
HDI: Highest density interval
HDPI: Highest density prediction interval
DICOM: Digital Imaging and Communications in Medicine

## DATA AND SUPPLEMENTAL MATERIALS

All data, code, and supplementary materials are available on the Open Science Framework project page (https://osf.io/g3vbe/).

## ACKNOWLEDGMENTS

We would like to thank the participants and FAU muscle physiology lab team members including but not limited to David Diaz, Shawn Dinh, Ethan Elkins, Caitlyn M. Meehan, Michael Morgan, Laura C. Canteri, Matthew Pritchard, and Chrisitan T. Macarilla for their assistance with data collection. Without these individuals, this project would not have been possible.

## AUTHOR CONTRIBUTIONS

Zac P. Robinson conceptualized the study, collected and analyzed all outcome measures, performed the statistical analysis, and drafted the manuscript. Michael C. Zourdos, James Steele, Eric R. Helms, Eric T. Trexler, Michael E. Hall, and Chun-Jung Huang all assisted in study design and edited the manuscript. James Steele also assisted in planning the statistical analysis. All other authors were an integral part of data collection, data entry, and edited the manuscript.

